# The RNA m6A reader YTHDF2 depletion phenocopies high glucose induced-cellular stress response in diabetic kidney disease

**DOI:** 10.64898/2026.10.23.761296

**Authors:** Harsha Sankar S H, Akhilesh Puranik, Kaustuv Tripathy, Nivedita Singh, Anuradha Kesharwani, Vipan Kumar Parihar, R Selvi Bharathavikru

## Abstract

RNA modification, especially the m6A RNA methylation, has been increasingly associated with several physiological and pathophysiological processes. The proteins that help in establishment of the modification and its removal along with the reader proteins that facilitate the functional outcome, together play an important regulatory role in disease manifestation. Diabetic kidney disease (DKD) is one such example, which is categorized as a multifactorial disorder that involves complex metabolic, oxidative, inflammatory, and fibrotic alterations, mediated through genetic, epigenetic and signalling cascades. However, the contribution of m6A-associated RNA regulators in DKD manifestation is relatively less explored. In this study, using HFD+STZ-induced diabetic mice and high glucose-treated renal cell models, we identified the m6A reader protein, Ythdf2 as a consistently downregulated candidate. YTHDF2 depletion in the renal cell line altered cellular viability, proliferation and increased oxidative stress under high-glucose conditions. Click-IT EU labeling further revealed altered RNA processing effects following YTHDF2 depletion and glucose stress. In corroboration with the above observations in animal and cell line models, independent human DKD transcriptomic datasets demonstrated conserved inflammatory and fibrotic signatures as well as reduced YTHDF2 expression in DKD glomeruli. The YTHDF2 downregulation was found to be inversely correlated with several disease-associated inflammatory and fibrotic genes. Together, these findings identify YTHDF2 dysregulation as a potential post-transcriptional mechanism associated with cellular stress and disease-associated molecular programs in DKD.

## Introduction

Diabetic kidney disease (DKD) is a leading cause of chronic kidney disease (CKD) and end-stage renal disease (ESRD). DKD affects 30% of type-I diabetic patients and 40% of people with type-II diabetes [1]. Over time, poorly controlled diabetes can damage the blood vessels in the kidneys that filter waste from the blood. This can lead to kidney damage and high blood pressure, which in turn exacerbates kidney damage by increasing pressure in the kidneys’ filtering system [2] [3]. Patients with DKD often require lifelong dialysis or kidney transplantation, causing a severe economic and social burden [4]. DKD is characterised by progressive reduction in glomerular filtration rate and elevated albuminuria on clinical assessment. The pathological characteristics of DKD include hyperglycemia-induced metabolic changes, glomerular basement membrane thickening, podocyte apoptosis, glomerular hypertrophy, mesangial expansion, tubular inflammation, oxidative stress, and renal fibrosis [5]. These significant characteristics of DKD are, in turn, driven by several molecular pathways, including TGF-β/Smad, which is the master regulator of renal fibrosis; AGE (Advanced glycation end products), which elevates ROS; and NF-κB, which in turn activates inflammatory cytokine production such as TNF-α, IL-1β, and IL-6 [6–7]. Beyond molecular pathways, epigenetic changes associated with DKD have been shown to contribute to metabolic memory, in which kidney injury persists even after improved glucose control [8]. Among the epigenetic mechanisms, histone modifications, DNA methylation, microRNAs, long non-coding RNAs, and m6A RNA methylation has been shown to contribute to the regulation of gene expression and the pathogenesis of DKD [9–10].

N6-methyladenosine (m6A) is the most abundant and extensively studied internal modification of eukaryotic mRNA and is also present in several non-coding RNAs, including lncRNAs, circRNAs, and microRNA precursors [11]. Unlike DNA methylation, m6A modification is a dynamic and reversible epitranscriptomic modification that enables rapid regulation of RNA metabolism in response to developmental, physiological, and pathological stimuli [12–13]. It is enriched around stop codons, within 3′ untranslated regions (3′UTRs), and in long internal exons, where it influences RNA fate through interactions with m6A RNA-binding proteins. The deposition, removal, and recognition of m6A are mediated by three classes of regulators—writers, erasers, and readers [14–15]. The writer complex is primarily composed of the catalytic subunit METTL3 and the RNA-binding scaffold METTL14, with accessory factors including WTAP, VIRMA, RBM15/RBM15B, ZC3H13, and HAKAI contributing to substrate recognition, localization, and methylation efficiency [16–17]. The erasers FTO and ALKBH5 remove m6A through oxidative demethylation [18–20]. Among the readers, the YTH-domain family including YTHDC1, YTHDF1, YTHDF2, and YTHDF3 regulates diverse aspects of RNA processing, translation, and degradation, with YTHDF2 being particularly associated with m6A-dependent transcript turnover [21–23]. Other m6A-associated readers include IGF2BPs, HNRNPA2B1/HNRNPG, and HuR, which contribute to RNA stability, processing, and post-transcriptional regulation. Collectively, these regulators enable m6A to fine-tune gene expression and RNA fate without altering the underlying nucleotide sequence, thereby facilitating cellular adaptation to developmental cues, environmental stress, and metabolic changes [24].

Emerging evidence indicates that m6A-associated RNA regulation contributes to cellular responses to hyperglycemic and diabetic renal stress, with alterations in m6A writers, erasers, and readers implicated in inflammation, oxidative stress, mitochondrial dysfunction, and renal injury [14, 25–28]. Individual studies have demonstrated cell and target-specific functions of m6A regulators in renal injury. For example, METTL3 promotes podocyte injury through m6A-dependent regulation of TIMP2 [29], while the METTL3–YTHDF2 axis regulates PINK1 and mitophagy in high-glucose-treated renal tubular cells [30]. METTL14 has been implicated in glomerular endothelial injury through regulation of α-Klotho [31], whereas WTAP/METTL14-dependent regulation of DKK3 and FTO-mediated regulation of NLRP3 have been associated with tubular injury and inflammatory responses, respectively [32–33]. Recent transcriptomic analyses have further highlighted differential expression of m6A-associated regulators across human DKD tissues and disease stages [34]. Despite these observations, the effects of hyperglycemic stress on m6A-associated RNA regulation across distinct renal cell types and developmental context remain poorly understood. In particular, comparatively little is known about the direct contribution of individual m6A readers to renal cellular responses to metabolic stress. Existing studies involving YTHDF2 in diabetic renal injury have largely examined its role within specific writer–reader pathways, including the METTL3–YTHDF2 axis, rather than addressing how YTHDF2 itself responds to hyperglycemic stress or how its depletion broadly affects renal cellular adaptation [30]. Thus, reader-specific mechanisms linking glucose stress to changes in RNA fate remain incompletely defined.

To address this question, we used M15 cells, derived from embryonic mouse mesonephric mesenchyme and representing an early nephron progenitor-like developmental context, to investigate cell-intrinsic responses to metabolic stress [35]. This developmental context is particularly relevant because metabolic and environmental cues during early kidney development can influence nephron formation and may have long-term consequences for renal function and disease susceptibility. Recent studies demonstrating that RNA transmethylation and METTL3-dependent pathways regulate nephron progenitor fate and nephrogenic potential further support a role for m6A-associated RNA regulation in kidney development [36]. Thus, examining how hyperglycemic stress alters m6A-associated RNA regulation in an early renal developmental context may provide insight into cellular responses relevant to DKD and into how adverse metabolic environments during kidney development could influence later renal susceptibility.

In this study, we therefore examined the expression and regulation of m6A-associated factors across distinct renal cellular contexts, including early developmental renal cells and differentiated inner medullary collecting duct derived-IMCD3 cell line [37] and investigated their response to hyperglycemic stress. YTHDF2 was of particular interest because of its established role in m6A-dependent RNA turnover and post-transcriptional control of transcript abundance. We subsequently focused on YTHDF2 to determine how its depletion influences renal cellular viability, proliferation, cell-cycle progression, oxidative stress and nascent RNA-associated activity. Finally, analysis of independent human DKD transcriptomic datasets was used to assess the relevance of YTHDF2-associated changes to inflammatory and fibrotic molecular signatures in human disease. Together, this approach provides a cell-contextual framework for understanding how m6A-associated RNA regulation may influence renal cellular adaptation to metabolic stress and its potential relevance to metabolic stress and establishes YTHDF2 as a potential molecular link between metabolic stress and renal cellular dysfunction, highlighting its possible role in the pathogenesis of metabolic renal stress related disorders like DKD.

## Materials and Methods

### Cell culture and treatment

M15 mouse mesonephric cells and IMCD3 inner medullary collecting duct-derived cells were maintained in DMEM (Thermo Fisher Scientific; 31600034) supplemented with 10% fetal bovine serum (FBS; HiMedia; RM9955) and 1% penicillin–streptomycin (Thermo Fisher Scientific; 15140122) at 37°C in a humidified atmosphere containing 5% CO₂.

For experimental treatments, cells were maintained under low-serum conditions (1% FBS) for 48 h. Normal glucose (NG; 5.5 mM glucose) was used as the control condition, whereas hyperglycemic stress was induced using 30 mM glucose (HG; Sigma-Aldrich; G7021). An osmotic control (HM) was established using 5.5 mM glucose supplemented with 24.5 mM mannitol (Sigma-Aldrich; M4125). NG, HG, and HM groups were maintained under identical serum concentrations and treatment durations.

### CRISPR-Cas9-mediated generation of YTHDF2-knockdown cells

Single-guide RNAs (sgRNAs) targeting YTHDF2 were designed using CHOPCHOP [38], and the sequences are provided in **Supplementary Table 1**. The selected sgRNAs were annealed using T4 polynucleotide kinase (NEB; M0201L) and cloned into the LentiCRISPR-v2 vector following BsmBI-v2 (NEB; R0739S) restriction digestion. Recombinant plasmids were verified by Sanger sequencing. Confirmed plasmids were transfected into HEK293T cells together with the required packaging and envelope vectors for lentiviral production. The resulting viral supernatants were used to transduce M15 cells. Transduced cells were selected with puromycin (InvivoGen; ant-pr-1; 2 μg/mL) for 14–28 days based on a previously established puromycin kill curve. YTHDF2 depletion was subsequently confirmed by western blotting.

### High fat diet (HFD) + Streptozotocin (STZ)-induced type 2 diabetic mouse model

Six-week-old male C57BL/6J mice were obtained from the NIH and maintained under standard housing conditions in the Animal House Facility at NIPER, Hajipur. The experimental protocol was performed as previously described by Singh et al. [39–40]. Mice were divided into control (NCD + vehicle) and diabetic (HFD + STZ) groups. Control animals were maintained on a normal-calorie diet containing 11.32% fat, 22.48% protein, and 63.0% carbohydrate, whereas the diabetic group received a high-fat diet containing 49.0% fat, 24.20% protein, and 29.23% carbohydrate. The HFD group additionally received two low doses of streptozotocin (30 mg/kg, intraperitoneally) on days 60 and 63 following initiation of dietary modification. Diabetes was confirmed by biochemical assessment of blood parameters as previously reported [39–40]. Following confirmation of diabetes, one cohort of animals was saline-perfused, and kidneys were collected for subsequent RT-qPCR analysis.

### RNA isolation and DNase treatment

Flash-frozen kidney tissues were collected from six animals per group, sectioned, weighed, and incubated with an equal volume of TrypLE™ Express Enzyme (1×; Thermo Fisher Scientific; 12604021) at 37°C for 5 min. The tissues were subsequently homogenized and centrifuged at 5,000 rpm for 5 min. The resulting pellet and supernatant were processed using TRIzol reagent (Thermo Fisher Scientific; 15596026), followed by chloroform (Sigma-Aldrich; C2432) extraction and centrifugation at 12,500 rpm for 15 min. The aqueous phase was collected in the presence of 25 μg linear polyacrylamide (LPA; Sigma-Aldrich; 56575-1ML), mixed with an equal volume of isopropanol (Sigma-Aldrich; 190764), and incubated for 10 min. RNA was precipitated by centrifugation at 12,500 rpm for 15 min at 4°C, washed twice with 70% ethanol, air-dried, and resuspended in 20–30 μL nuclease-free water. RNA concentration and purity were assessed using a NanoDrop spectrophotometer, and RNA integrity was evaluated by agarose gel electrophoresis. RNA isolation from cell pellets was performed using the same procedure beginning with TRIzol treatment. For DNase treatment, RNA was incubated with RNase-Free DNase (QIAGEN; 79254) and the corresponding reaction buffer according to the manufacturer’s instructions.

### cDNA synthesis and RT-qPCR

Following assessment of RNA integrity, 1 μg of total RNA was used for cDNA synthesis with 1 μL oligo(dT) or random primers. RNA and primers were incubated at 70°C for 5 min and subsequently cooled on ice for 5 min. cDNA was synthesized in a 20 μL reaction using the GoScript™ Reverse Transcriptase cDNA Synthesis Kit (Promega; A5001) according to the manufacturer’s instructions. The resulting cDNA was diluted 1:10 before RT-qPCR.

RT-qPCR was performed using PowerUp™ SYBR™ Green Master Mix (Thermo Fisher Scientific; A25742) with 400 nM each of forward and reverse primers and 1 μL diluted cDNA per reaction. Primer sequences are provided in **Supplementary Table 2.**

### Western blotting

Total protein was extracted from renal cells using RIPA buffer supplemented with a protease inhibitor cocktail (HiMedia; ML051). Protein concentration was determined using the Bradford assay, and equal concentration of protein was separated by SDS-PAGE and transferred onto PVDF membranes (Bio-Rad; 1620177) using a semi-dry transfer method.

Membrane was blocked with 5% skimmed milk (Himedia; GRM1254) for 1 h at room temperature and incubated overnight at 4°C with anti-YTHDF2 antibody (ABclonal; A28394; 1:5,000) and β-actin HRP-conjugated antibody (Abclonal, AC028; 1:50,000), whereas YTHDF2 was detected using a primary antibody followed by an HRP-conjugated anti-rabbit secondary antibody (CST, 7074P2; 1:2,000). Following secondary-antibody incubation, membranes were washed three times with 1× TBST. Protein signals were visualized using Clarity ECL substrate (Bio-Rad; 1705061) and quantified using ImageJ.

### Cell proliferation assay

For proliferation analysis, 10,000 cells were seeded in 24-well plates and allowed to attach overnight. Cells were subsequently exposed to HM or HG under low-serum conditions for 48 h. Cells were harvested by trypsinization, and cell numbers were determined using a CASY cell counter.

### Cell viability assay

Cell viability was assessed using the MTT assay. Briefly, 2,500 cells were seeded in 96-well plates and allowed to attach overnight before exposure to HM or HG under low-serum conditions. At the indicated time points, 10 μL MTT reagent (HiMedia; TC191; 5 mg/mL) was added to each well and incubated for 4 h at 37°C. Formazan crystals were dissolved in DMSO (SRL; 28580), and absorbance was measured at 570 nm using Tecan Infinite® M Plex microplate reader. Cell viability was expressed relative to the corresponding control using background-corrected absorbance values.

### Intracellular ROS measurement

Intracellular reactive oxygen species (ROS) levels were measured using DCF-DA (Sigma-Aldrich; D6883). A total of 2,500 cells were seeded in black 96-well plates and maintained overnight before exposure to HM or HG under low-serum conditions for 48 h. H₂O₂-treated cells (10 mM for 30 min) were included as a positive control. Following treatment, cells were washed with PBS and incubated with 5 μM DCF-DA for 30 min at 37°C in the dark. Fluorescence was measured using a TECAN Infinite® M Plex microplate reader at approximately 485 nm excitation and 535 nm emission wavelengths.

### Cell-cycle analysis

For cell-cycle analysis, cells were harvested at approximately 80% confluence from 90-mm culture dishes and washed with 1× PBS. Cells were fixed in 70% ethanol for 30 min at 4°C and centrifuged at 2,000 rpm for 3 min. Following two washes with PBS, cells were stained with RNase (100 μg/mL) and propidium iodide (HiMedia; ML067; 50 μg/mL) in 200 μL PBS for 10 min in the dark. DNA content was analyzed by Beckman Coulter CytoFLEX LX flow cytometer using the Y585-A channel.

### EU pulse-chase labeling and Click-iT detection of nascent RNA

Nascent RNA synthesis was assessed using 5-ethynyl uridine (EU) metabolic labeling followed by Click-iT detection. A total of 10,000 cells stably expressing empty vector (EV) or YTHDF2-knockdown constructs (KD1 and KD2) were seeded on poly-L-lysine-coated coverslips and maintained under NG, HG, or HM conditions for 48 h. Nascent RNA was labeled with 1mM EU (Thermo Fisher Scientific; E10345) for 15 min at 37°C. Following labeling, the EU-containing medium was replaced with NG medium. For transcriptional inhibition, cells were subsequently treated with actinomycin D (HiMedia; TC018; 25 μg/mL) for 2 h. Untreated cells and cells exposed to actinomycin D before EU labeling were included as negative controls.

Cells were fixed with 4% paraformaldehyde for 10 min at room temperature and permeabilized with 0.25–0.3% Triton X-100 (Sigma-Aldrich; T8787) for 10 min. Following blocking with 2% BSA, incorporated EU was detected using an Alexa Fluor 555 azide-based Click-iT reaction (Thermo Fisher Scientific; A20012). Coverslips were counterstained with DAPI (Sigma-Aldrich; 11196301) and mounted for imaging. Confocal images were acquired using a Leica STELLARIS 8 FALCON confocal microscope at the Central Advanced Instrumentation Facility (CAIF), IISER Berhampur. EU-associated fluorescence was quantified using ImageJ. Total fluorescence intensity was measured per cell and normalized to the corresponding EV control mean fluorescence intensity within each biological replicate. For actinomycin D-treated samples, residual EU-associated fluorescence was quantified following transcriptional inhibition and normalized to the corresponding initial EU-associated signal and EV control.

### Transcriptomic and proteomic meta-analysis

Differential expression data from mouse kidney transcriptomics was obtained from a previously published RNA-seq study of diet or streptozotocin-induced diabetic nephropathy [39]. Differential expression of the genes listed in **Supplementary Table 3** was assessed using NOISeqBIO, with log₂ fold change (log₂FC) and posterior probability of differential expression [prob(DE)] used for statistical evaluation. Genes with log₂FC ≥ 1.0 and prob(DE) < 0.05 were considered differentially expressed.

Differential protein abundance was assessed using the corresponding published proteomic dataset [39] and Proteome Discoverer output. Proteins with an absolute log₂FC ≥ 1.0 and Benjamini–Hochberg (BH)-adjusted p < 0.05 were considered differentially expressed. Gene Ontology biological process enrichment analysis was performed using clusterProfiler::enrichGO, and enriched terms were visualized as dot plots.

### Analysis of human DKD transcriptomic datasets

Three publicly available human DKD transcriptomic datasets were obtained from the Gene Expression Omnibus (GEO) using the GEOquery package: GSE142025, comprising kidney biopsy transcriptomes generated by high-throughput RNA sequencing on the Illumina HiSeq 4000 platform; GSE96804, comprising microdissected glomerular samples profiled using the Affymetrix Human Transcriptome Array 2.0; and GSE104954, comprising tubulointerstitial transcriptomes generated using Affymetrix Human Genome U133 Plus 2.0 and Human Genome U133A Array platforms [41–43]. GSE142025 included 28 diabetic nephropathy and 9 normal kidney samples, while GSE96804 comprised 41 diabetic nephropathy and 20 control glomerular samples. GSE104954 contains multiple kidney disease groups and healthy living-donor samples; for the present analysis, the diabetic nephropathy subset of 17 samples and 21 healthy living-donor samples was selected. Thus, the three datasets provided complementary transcriptomic profiles of diabetic kidney disease across kidney biopsy, glomerular, and tubulointerstitial compartments.

Samples were categorised as healthy controls or DKD irrespective of disease stage. Differential expression analysis was performed using the limma workflow. A design matrix without an intercept was fitted using lmFit, followed by extraction of the DKD-versus-control contrast using contrasts.fit. Moderated t-statistics were calculated using empirical Bayes shrinkage with eBayes, and p-values were adjusted using the Benjamini–Hochberg method. Probe-to-gene mapping was performed using the respective platform annotations. For GSE142025, per-sample expression files were obtained from GEO, merged into a single expression matrix, and analysed using the same limma-based workflow.

Genes significantly upregulated (p < 0.05) across the three datasets were identified using a three-way Venn analysis. Genes commonly upregulated across all three renal compartments were subsequently visualised using lollipop plots to compare their expression changes across datasets.

Genes of interest were classified into predefined biological categories associated with DKD progression, including fibrosis, inflammation, oxidative stress, cellular stress and m6A-associated regulators. The same categorisation was applied across datasets for consistency. YTHDF2 log₂FC values were compared with those of selected DKD-associated genes, and Pearson correlation analysis was performed using the cor.test() function in R.

### Statistical analysis

Statistical analyses were performed using GraphPad Prism (GraphPad Software). Comparisons between two independent groups were performed using an unpaired, two-tailed Student’s t-test. Cell culture experiments were performed using three independent biological replicates (n = 3), while mouse experiments included five or six biological replicates per group (n = 5–6), as indicated in the respective figure legends. Data are presented as mean ± SD, and p < 0.05 was considered statistically significant. Statistical significance was indicated as follows: p < 0.05, p < 0.01, p < 0.001, and p < 0.0001

All bioinformatic analyses were performed in R (v4.4.2) using RStudio (v2026.04.0+526), with ggplot2 used for data visualization.

## RESULTS

### Diabetic kidney disease induction via HFD and STZ alters the expression of m⁶A modifying machinery

Diabetic kidney disease (DKD) is characterized by well-established molecular hallmarks, including inflammation, oxidative stress, and extracellular matrix (ECM) remodeling and fibrosis. We first examined whether these disease-associated molecular signatures were evident in the HFD+STZ-induced diabetic kidney model. To this end, we performed a targeted reanalysis of previously generated transcriptomic data from the study reported by Singh *et al.* 2025 [39]. Predefined panels of genes associated with cellular stress, inflammation, and fibrotic remodeling were curated, and their expression patterns were evaluated in the renal transcriptomic dataset. This analysis revealed substantial alterations across all three disease-relevant categories **(Figure 1A)**. Within the cellular stress-associated gene panel, *Hmox1, Cdkn1a,* and *Nfe2l2* were upregulated, whereas *Casp9* and *Bax* were downregulated in diabetic kidneys. Among inflammation-associated genes, *Casp8, Cxcl10,* and *Nfkbia* showed increased expression, whereas *Rela* and *Il33* were decreased. The fibrotic markers *Col3a1*, *Mmp13*, *Pdgfrb*, *Mmp12*, and *Tgfbr1* were upregulated, whereas *Col1a2*, *Tgfb2*, *Acta2*, *Tgfb1*, and *Ccn2* showed reduced expression. These findings demonstrate substantial alterations in molecular signatures associated with cellular stress, inflammation, and fibrotic remodeling in the HFD+STZ-induced diabetic kidney.

**Figure 1:**
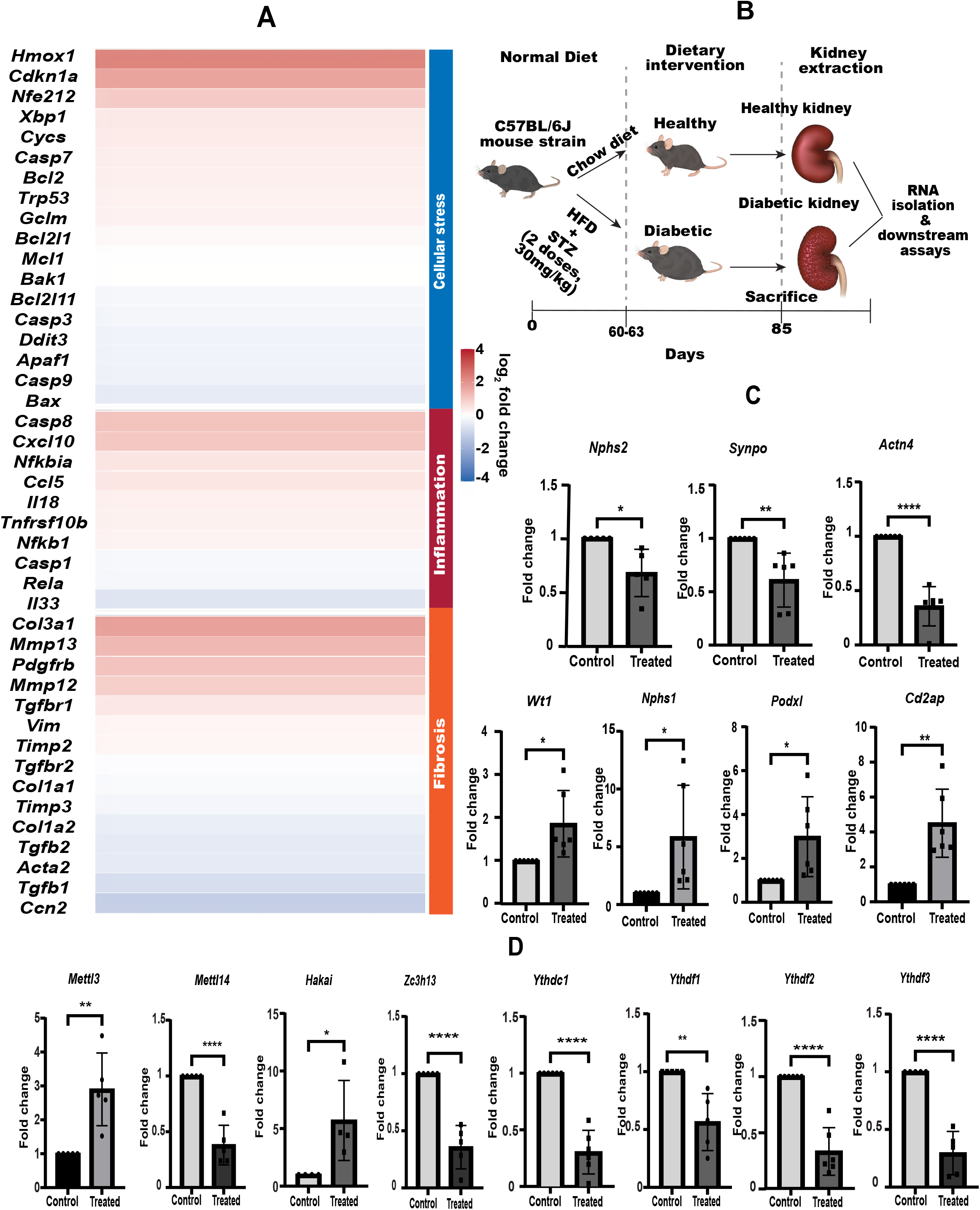
**A.** Heatmap representation of meta-analysis of differential expression of key genes involved in inflammation, apoptosis and stress responses in diseased samples (HFD/STZ treated mice) relative to healthy samples (NCD fed mice). The vertical axis denotes the change in log2FC values, and the horizontal rows denote the genes involved in disease progression. Colour intensity reflects relative expression levels (scaled values), with red indicating higher expression and blue indicating lower expression. **B**. Schematic representing experimental workflow for RT-qPCR-based gene expression analysis in the HFD/STZ-induced diabetic nephropathy mouse model. **C.** Bar plot displaying relative expression of significantly dysregulated podocyte-associated markers in diseased samples compared to healthy controls. **D.** Bar plot displaying relative expression of significantly dysregulated m6A writers and readers in diseased samples compared to healthy controls. Bars represent mean +/− SD. Statistical significance determined by Student’s t-test. (*p < 0.05, * *p < 0.01, * * *p < 0.001 and * * * *p < 0.0001) N=5 for genes *Nphs2*, *Mettl3*, *Mettl14*, *Hakai*, *Zc3h13*, *Ythdf1* and *Ythdf3* and N=6 for genes *Synpo*, *Actn4*, *Wt1*, *Nphs1*, *Podxl*, *Cd2ap*, *Ythdc1* and *Ythdf2* (N represent independent biological replicates).

In addition to the targeted analysis, we performed Gene Ontology (GO) enrichment analysis using the whole renal transcriptomic dataset to identify broader biological processes associated with diabetic stress. This analysis revealed significant enrichment of biological processes related to extracellular matrix organization, extracellular structure organization, angiogenesis, organic hydroxy compound metabolism, organic acid transport, and carboxylic acid transport, among others **(Supplementary Figure** - **S1A)**. The enrichment of ECM and extracellular structure-associated processes is consistent with the well-established role of extracellular matrix remodeling in diabetic renal injury. Furthermore, enrichment of pathways related to angiogenesis and organic acid transport suggests alterations in renal vascular and metabolic processes, together with potential perturbation of tubular transport functions. Collectively, these transcriptome-wide pathway alterations support the presence of extensive structural, metabolic, and transport-associated changes characteristic of diabetic kidney injury in the HFD+STZ model.

Although the targeted re-analysis provided evidence of alterations in disease-associated inflammatory, cellular stress, and fibrotic signatures, it did not reveal significant changes in the selected podocyte-associated genes or m6A regulatory factors within the transcriptomic dataset. Given that podocyte injury is a prominent feature of DKD and that altered m6A-associated RNA regulation was the primary focus of this study, we therefore directly examined the expression of podocyte-associated genes and m6A regulatory factors in HFD+STZ kidney samples by RT-qPCR, following the experimental workflow shown in **Figure 1B**.

Analysis of podocyte-associated genes revealed significant alterations in multiple components of the podocyte molecular signature **(Figure 1C)**. Expression of the slit diaphragm-associated genes podocin *(Nphs2),* synaptopodin *(Synpo),* and α-actinin-4 *(Actn4)* was significantly reduced in diabetic kidneys, whereas wt1 *(Wt1),* nephrin *(Nphs1),* podocalyxin *(Pdxl),* and cd2ap (*Cd2ap)* showed significantly increased expression. These changes indicate substantial perturbation of the podocyte molecular phenotype in HFD+STZ-induced diabetic kidneys and are consistent with podocyte injury and adaptive remodeling under diabetic conditions.

To determine whether the alterations in podocyte-associated molecular signatures were also reflected at the protein level, we performed a targeted analysis of previously generated proteomic data from the same animal cohort. Analysis of selected podocyte-associated proteins revealed significant alterations in the diabetic kidneys, including reduced expression of α-actinin-4 (ACTN4) and nephrin (NPHS1), consistent with the changes observed in podocyte-associated transcripts. We further examined predefined proteins associated with cellular stress, renal fibrosis, and inflammation and identified significant alterations in several proteins involved in oxidative stress. Among these, TXN2, GSS, GCLM, and PRDX3 showed increased expression, whereas IDH2, GSTM5, and GLS were decreased in diabetic kidneys **(Supplementary Figure** - **S1B)**. To assess broader pathway-level changes, GO enrichment analysis was subsequently performed using the whole proteomic dataset. This analysis revealed major alterations in biological processes associated with oxidative stress responses, redox-related metabolism, and cellular responses to oxidative stress **(Supplementary Figure** - **S1C)**. Collectively, these findings demonstrate that the HFD+STZ-induced diabetic kidney exhibits alterations in podocyte-associated proteins together with pronounced proteome-wide changes related to oxidative stress, further supporting the presence of substantial cellular and molecular perturbations in this DKD model.

Several studies have reported alterations in epigenetic regulators in DKD; however, comparatively less is known about changes in the epitranscriptomic machinery in diabetic renal injury. We therefore examined the expression of m6A-associated regulatory factors, given our primary interest in understanding the contribution of m6A-associated RNA regulation to DKD. RT-qPCR analysis revealed broad alterations in the expression of components of the m6A regulatory machinery **(Supplementary Figure** - **S1D**). Among the m6A-associated factors examined, the catalytic methyltransferase *Mettl3* showed significantly increased expression, consistent with previous reports implicating METTL3 in diabetic kidney injury [31]. *Hakai* was also significantly increased, whereas components of the methyltransferase-associated complex, including *Mettl14* and *Zc3h13*, showed significantly reduced expression compared with control kidneys. Notably, several YTH-domain-containing m6A readers, including *Ythdc1, Ythdf1, Ythdf2,* and *Ythdf3*, were significantly downregulated in diabetic kidney tissue **(Figure 1D)**. These findings indicate that the m6A regulatory machinery undergoes substantial remodeling in the HFD+STZ-induced DKD model and prompted us to investigate the molecular mechanism of these m6A-associated factors under hyperglycemia conditions in renal cell models.

### High glucose induced differential alteration of m6A regulatory factors in renal cell line models

To determine whether the dysregulation of the m6A regulatory machinery observed in HFD+STZ-induced diabetic kidneys was recapitulated under hyperglycemic stress in renal cell models, we examined the expression of m6A-associated factors in two renal cell lines: M15 cells, derived from the early mesonephric developmental stage of the kidney, and IMCD3 cells, derived from the inner medullary collecting duct. As the expression profile of the broader m6A regulatory machinery has not been extensively characterised in these renal cell line models, we first examined the basal expression of a panel of m6A writers, erasers, and readers by RT-qPCR. All m6A-associated factors that were tested, were detectable in both M15 and IMCD3 cells **(Supplementary Figure** - **S2A),** indicating the presence of a functional repertoire of m6A regulatory components in both renal cell line models. Comparison of the basal expression profiles between M15 and IMCD3 cells revealed differential expression of several m6A regulatory factors. The writer-associated factors *Rbm15*, *Rbm15b* and the reader *Ythdf3* showed an increased expression in IMCD3 cells, whereas *Wtap*, *Ythdf1*, *Ythdf2*, *Hnrnpa2b1*, *Igf2bp1*, and *Igf2bp3* showed decreased expression relative to M15 cells **(Supplementary Figure** - **S2B)**. These differences indicate that the expression landscape of m6A regulatory factors varies between renal cell lines representing distinct developmental compartmentalization of these modulators.

We next investigated whether exposure to high glucose alters the expression of m6A regulatory factors in M15 and IMCD3 cells that were treated with normal glucose (NG) or high glucose (HG) for 48 h, as indicated in the experimental workflow shown in **Figure 2A**. In M15 cells, high-glucose exposure resulted in broad alterations in the expression of m6A-associated factors **(Figure 2B)**. Among these, the writer-complex-associated factor *Virma* and the reader *Igf2bp3* were significantly upregulated, whereas *Mettl14*, *Rbm15*, *Ythdc1*, and *Ythdf2* were significantly reduced **(Figure 2C)**. These findings indicate that high-glucose exposure substantially perturbs the m6A regulatory landscape in M15 renal cells. We next examined the response of the m6A machinery to high glucose in IMCD3 cells. A distinct pattern of dysregulation was observed, with significant increased expression of several m6A readers, including *Ythdf1*, *Hnrnpa2b1*, *Hnrnpg*, *Igf2bp1*, and *Igf2bp3*, whereas multiple writer-associated factors, including *Mettl3*, *Mettl14*, *Rbm15*, *Rbm15b*, *Virma*, and *Zc3h13* showed a decreased expression in comparison to normal glucose (NG) treated samples. **(Figure 2D and Supplementary Figure** - **S2C)**. In addition, the readers *Ythdc1*, *Ythdf2*, and *Hur* were significantly reduced following high-glucose exposure. Thus, although high glucose broadly perturbed the m6A regulatory machinery in both renal cell lines tested, the specific pattern of dysregulation differed between M15 and IMCD3 cells, suggesting cell specific and context-dependent responses of the m6A regulatory network to metabolic stress.

**Figure 2:**
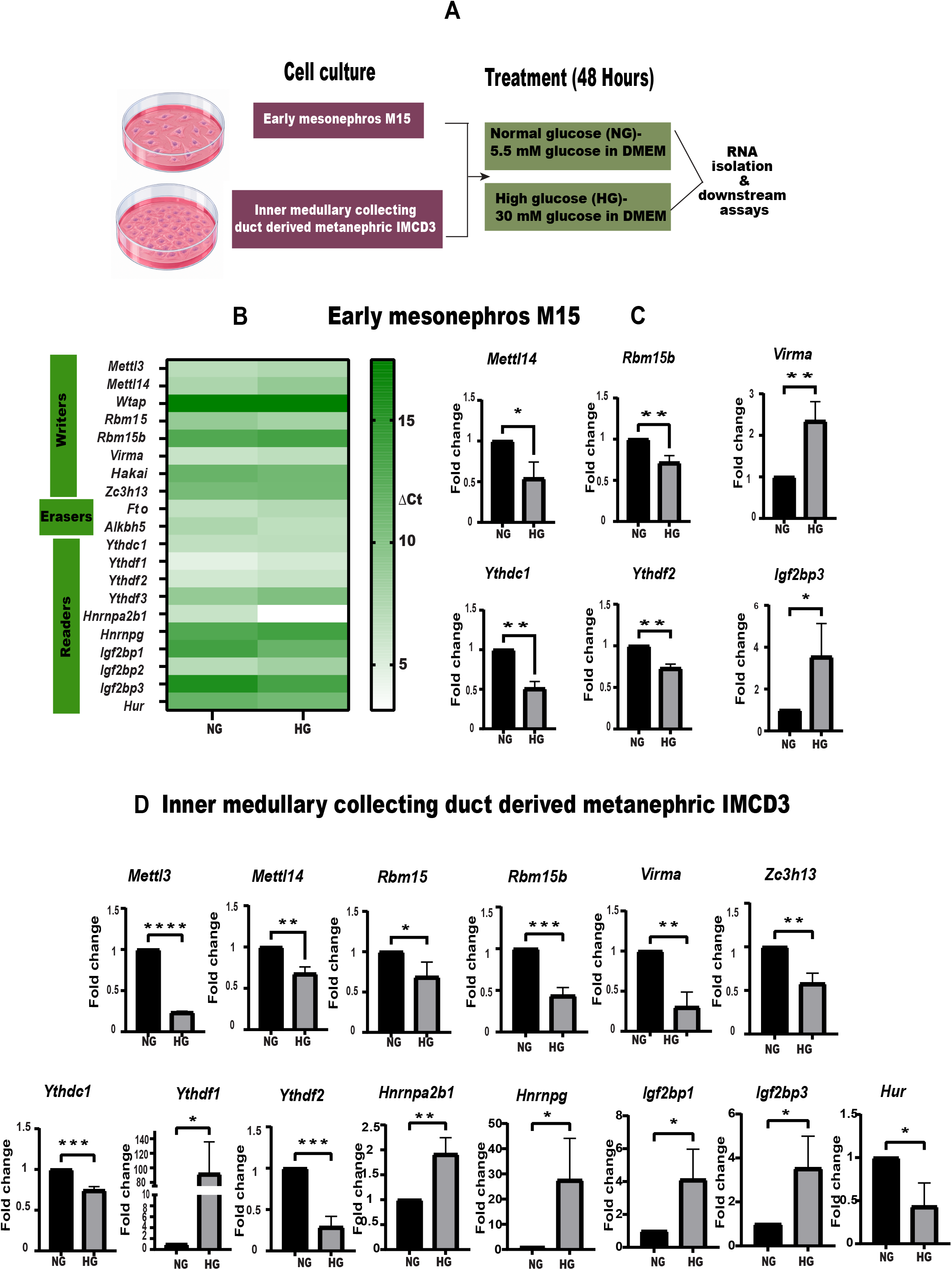
**A**. Schematic representing experimental workflow for RT-qPCR-based gene expression analysis of m6A regulators in the high glucose-treated mouse-derived renal cell lines, M15 (mesonephros-derived) and IMCD3 (inner medullary collecting duct-derived) cell lines. **B.** Heat map representing delta CT values of m6A regulators in high glucose (HG −30mM) treated M15 cell lines with respect to normal glucose treated cells (NG-5.5 mM). **C.** Bar plot showing relative expression of significantly dysregulated m6A writers and readers in HG-treated M15 cells compared to the NG-treated cells. **D.** Bar plot displaying the relative expression of significantly dysregulated m6A writers and readers in HG-treated IMCD3 cells compared to the NG-treatment. Data are presented as mean ± SD from N= 3 independent biological replicates. (* *p < 0.01, * * *p < 0.001 and ns indicates non-significant) Statistical significance determined by Student’s t-test.

Notably, writer *Mettl14* and readers *Ythdc1* and *Ythdf2* emerged as common m6A regulators showing consistent downregulation across the HFD+STZ kidney and high-glucose-treated renal cell models. Given the established role of YTHDF2 in m6A-dependent RNA turnover and decay, together with the relatively limited understanding of its contribution to glucose-induced renal cellular stress, we selected YTHDF2 for further mechanistic investigation.

To further determine whether the reduction in *Ythdf2* expression was specifically associated with glucose exposure rather than osmotic stress, *Ythdf2* transcript levels were independently assessed in M15 cells following NG, HM, and HG treatment. *Ythdf2* expression was significantly reduced under HG relative to both NG and HM conditions **(Supplementary Figure** - **S2D)**, further supporting a glucose-associated transcriptional response that cannot be attributed solely to increased osmolarity. Collectively, these findings demonstrate consistent downregulation of *Ythdf2* expression in high-glucose-treated renal cell models and HFD+STZ-treated mouse kidneys, prompting further investigation into the functional consequences and underlying molecular mechanisms of YTHDF2 depletion under hyperglycemic stress.

### YTHDF2 reduction promotes cellular stress responses phenocopying high-glucose exposure in M15 cells

To investigate the functional consequences of YTHDF2 depletion under hyperglycemic stress, we generated stable YTHDF2-depleted M15 renal cell lines using a LentiCRISPRv2-based CRISPR/Cas9 system. Two single-guide RNAs (sgRNAs), targeting exon 4 (sg1) and exon 5 (sg2) of *Ythdf2* **(Supplementary Figure** - **S3A)**, were cloned into the LentiCRISPRv2 vector, and the resulting constructs were validated by Sanger sequencing. Following lentiviral transduction and puromycin selection, stable M15 cell populations generated using sg1 and sg2 were designated KD1 and KD2, respectively, while an empty-vector-transduced M15 cell line (EV) was used as the control. Western blot analysis confirmed a reduction of more than 50% of YTHDF2 expression in both KD cells (KD1 and KD2) cells relative to EV cells, confirming effective YTHDF2 depletion **(Figure 3A, 3B)**. Interestingly, YTHDF2-depleted cells exhibited a visibly elongated morphology compared with EV control cells **(Supplementary Figure** - **S3B)**. To quantify these morphological changes, cellular aspect ratio (AR) and solidity were measured using ImageJ in EV, KD1, and KD2 cells under normal-glucose (NG), hyperosmotic-control (HM), and high-glucose (HG) conditions. This analysis was performed to determine whether the observed morphological alterations were associated primarily with YTHDF2 depletion rather than glucose treatment. YTHDF2-depleted cells exhibited an increased AR distribution and reduced solidity relative to EV cells across the treatment conditions **(Supplementary Figure** - **S3C and S3D)**. Together, these findings indicate a more elongated and less compact cellular morphology, suggesting that YTHDF2 depletion is associated with a mesenchymal-like cellular phenotype, irrespective of glucose treatment conditions.

**Figure 3:**
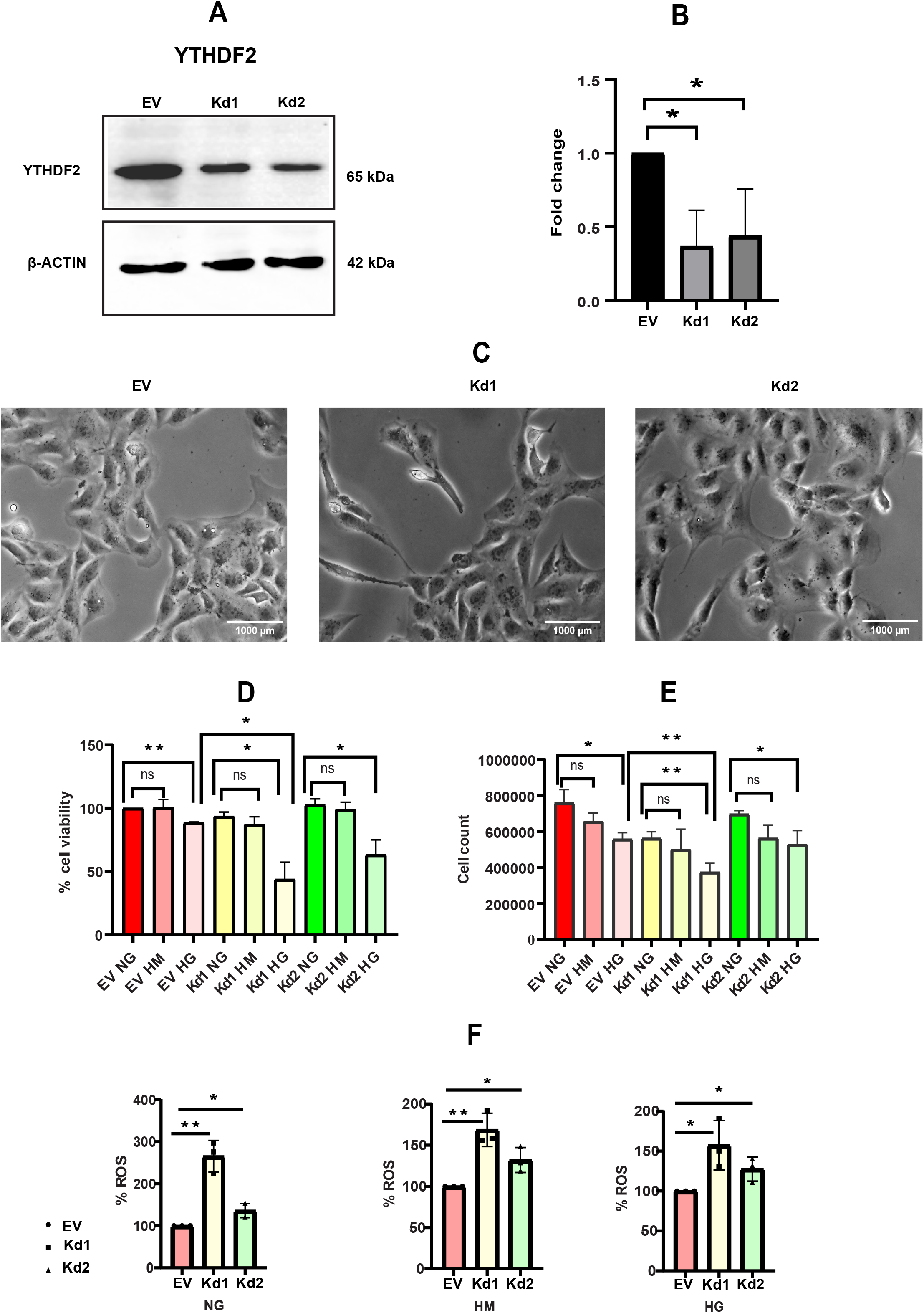
**A.** Western blot analysis of YTHDF2 expression in genome edited cell lines compared to ◻-actin as the loading control. **B.** The bar plot represents the quantification of relative protein levels in the empty vector control compared with the YTHDF2 KD1 and KD2 transduced cell lines. **C.** Bar plot showing the percentage cell viability in M15 EV cells in comparison with YTHDF2 KD cells across different glucose treatments (NG, HM and HG). Data normalized to M15 EV NG cells. **D.** Bar plot showing the cell count in M15 EV in comparison with YTHDF2 KD cells across different glucose treatments. **E.** Bar plot showing the percentage ROS production at different glucose treatment conditions. Each plot is normalised to the respective EV. (Data are presented as mean ± SD from *N* = 3 independent biological replicates. (*p < 0.05, * *p < 0.01, * * *p < 0.001 and * * * *p < 0.0001) Statistical significance determined by Student’s t-test).

We next examined whether YTHDF2 depletion influenced cellular cytotoxicity and proliferation under hyperglycemic stress. M15 cells (EV, KD1 and KD2) were exposed to NG, HM, or HG conditions, and cellular viability was assessed by MTT assay. HM treatment did not significantly alter viability relative to NG in any cell population, whereas HG exposure significantly reduced viability in all cells, confirming the induction of glucose-associated cellular cytotoxicity **(Figure 3C)**. Notably, KD1 cells exhibited a significant further reduction in viability compared with EV cells under HG conditions, whereas KD2 cells showed a similar trend of reduction in cell viability. These findings suggest that YTHDF2 depletion may increase cellular susceptibility to high-glucose-induced loss of viability. Consistent with the cell viability measurements, direct assessment of cell numbers using a cell counter revealed a similar pattern of reduced cellular proliferation under HG conditions, with YTHDF2-depleted cells exhibiting a further reduction relative to EV cells **(Figure 3D).**

To determine whether altered proliferation was associated with changes in cell-cycle progression, we performed propidium iodide-based flow cytometric analysis. Both KD cell populations exhibited an increased proportion of cells in S phase relative to control M15 EV cells, indicating altered cell-cycle progression following YTHDF2 depletion **(Supplementary Figure** - **S3E, S3F & S3G)**. Collectively, these findings indicate that YTHDF2 depletion alters cell viability and proliferation and is associated with altered cell-cycle progression, particularly under high-glucose stress.

Because oxidative stress is a well-established consequence of high-glucose exposure in renal cells, we next investigated whether YTHDF2 depletion affected intracellular reactive oxygen species (ROS) levels using DCFDA fluorescence under NG, HM, and HG conditions. Unstained cells were included as negative controls (NC), while H₂O₂-treated cells were used as positive controls (PC) to validate the ROS detection assay. As expected, NC cells exhibited minimal fluorescence compared with DCFDA-stained M15 EV cells under NG conditions, whereas H₂O₂-treated PC cells showed markedly higher fluorescence, confirming the functionality of the assay (**Supplementary Figure** - **S3H**). To determine whether YTHDF2 depletion was associated with altered oxidative stress, ROS levels in YTHDF2-depleted cells were compared with those in EV control cells under each glucose condition. The KD cells exhibited significantly higher ROS levels than EV cells across the treatment conditions (**Figure 3E**). Notably, this increase was observed irrespective of glucose condition, indicating that YTHDF2 depletion itself is associated with elevated intracellular ROS levels. These findings suggest that loss of YTHDF2 may contribute to increased basal oxidative stress and potentially influence the cellular response to oxidative challenges.

Altogether, these data demonstrate that YTHDF2 depletion impairs cell viability and proliferation, disrupts cell-cycle progression, and promotes intracellular ROS accumulation in M15 cells. YTHDF2-depleted cells also displayed altered morphology, with an elongated, mesenchymal-like appearance, indicating that YTHDF2 loss may affect cellular phenotype in addition to growth and oxidative-stress responses. Similarly, the effects of YTHDF2 depletion and high-glucose stress on cellular growth and oxidative stress suggests that YTHDF2 may contribute to the regulation of renal cellular adaptation to hyperglycemic conditions. Given the established role of YTHDF2 as an m6A RNA modification reader involved in post-transcriptional RNA regulation, we next investigated whether YTHDF2 depletion alters global RNA dynamics under normal and high-glucose conditions.

### YTHDF2 depletion perturbs nascent RNA synthesis and RNA turnover dynamics under high glucose conditions

To investigate whether YTHDF2 depletion alters global RNA dynamics under hyperglycemic conditions, we assessed nascent RNA synthesis using 5-ethynyl uridine (EU) metabolic labelling coupled with Click-iT chemistry. M15 EV, KD1, and KD2 cells were exposed to normal glucose (NG), hyperosmotic mannitol (HM), or high glucose (HG), followed by a short pulse of EU labelling. EU incorporation was subsequently visualised by conjugation with Alexa Fluor 555 azide, and the total cellular fluorescence intensity was quantified as a measure of nascent RNA synthesis **(Figure 4A)**. To establish the specificity of the Click-iT signal, two negative controls were included: cells processed in the absence of EU and cells treated with actinomycin D prior to EU labelling. Both controls showed minimal detectable fluorescence in the Alexa Fluor 555 dye corresponding red channel (HyD S3 detector), confirming that the observed signal was predominantly dependent on EU incorporation **(Supplementary Figures** - **S4A and S4B)**. Following EU labelling, the fluorescence signal was predominantly concentrated within nucleolar regions, consistent with the contribution of newly synthesised ribosomal RNA to the total nascent RNA signal **(Supplementary Figure** - **S4C, S4D, S4E)**. Quantification of EU fluorescence revealed a significant reduction in nascent RNA labelling following HG exposure in M15 cells compared with the corresponding NG and HM conditions **(Figure 4B**, **4C)**. These findings indicate that high-glucose stress is associated with reduced global nascent RNA synthesis in M15 cells. Notably, YTHDF2-depleted cells exhibited a marked reduction in EU incorporation compared with EV cells irrespective of glucose treatment condition **(Figure 4D)**, indicating that YTHDF2 depletion is associated with reduced global nascent RNA synthesis irrespective of glucose condition.

**Figure 4:**
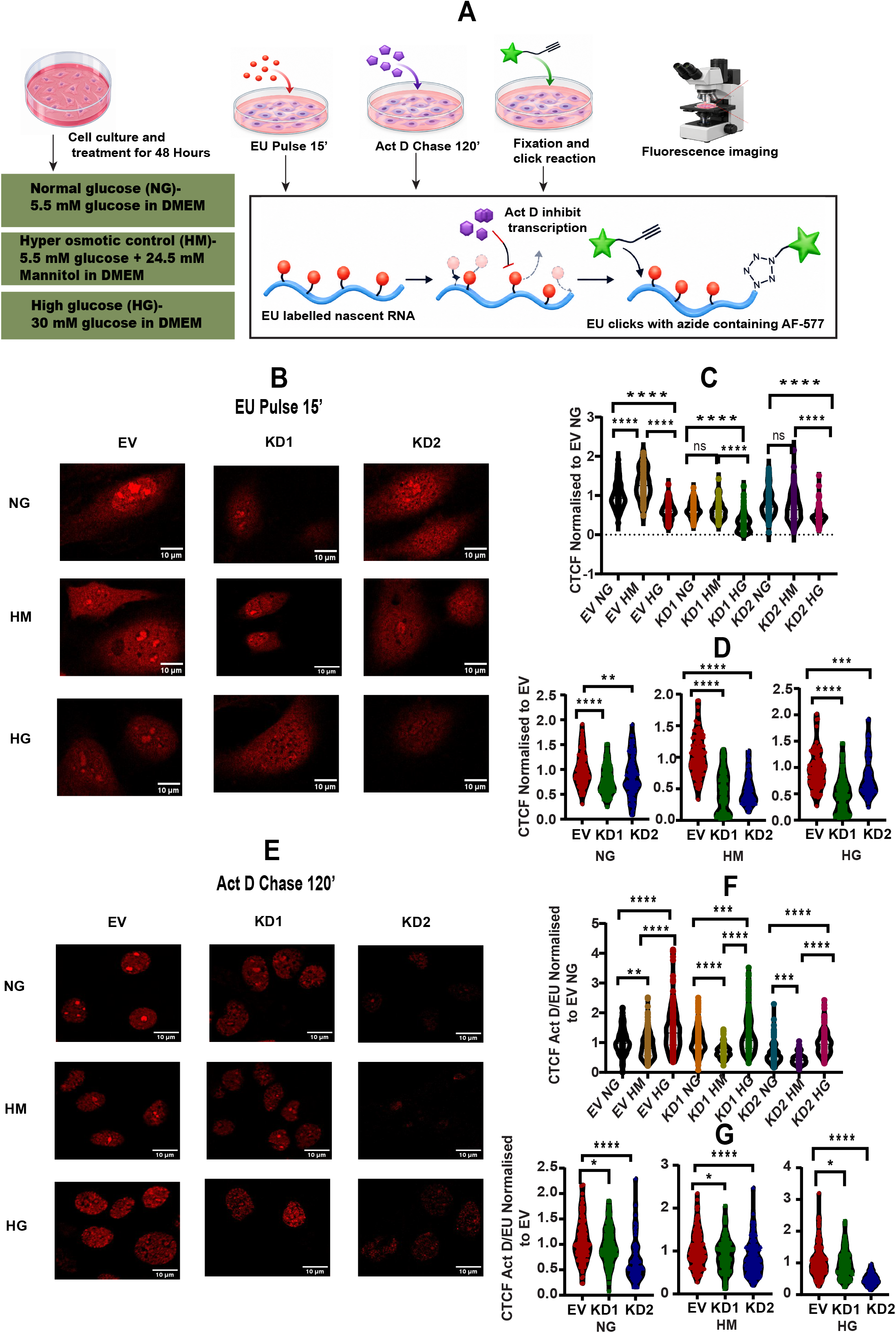
**A.** Schematic representing experimental workflow of click chemistry. **B.** Representative image showing AF-577 intensity depicting nascent RNA synthesis measured by EU (5-Ethynyl uridine) staining in YTHDF2 knockdown cells with respect to empty vector control across NG, HM and HG. **C.** Violin plot depicting the quantification of nascent synthesis of RNA in control (EV) and YTHDF2 knockdown (kd) cells across different treatment conditions (NG, HM and HG) (n= (50-100) *4). **D.** Violin plots showing the normalised intensity of EU labelled cells of YTHDF2 KD with respect to M15 EV in NG, HM and HG (n= (50-100) *4). **E.** Representative image showing AF-577 intensity of EU labelled cells after treatment with Actinomycin-D for 2 hours in YTHDF2 knockdown cells with respect to empty vector control across NG, HM and HG. **F.** Violin plot indicating quantification of RNA kinetics (intensity of act D/EU normalised to EV NG) in control (EV) Vs YTHDF2 knock down cells across different treatment conditions (50-100) *3). **G.** Violin plot showing the normalised intensity of Act D / EU to EV in YTHDF2 KD cells with respect to M15 EV across NG, HM and HG (50-100) *3). Data are presented as violin plots showing the distribution of individual observations, *N* = 3 independent biological replicates. CTCF denotes Corrected total cell fluorescence. (* *p < 0.01, * * *p < 0.001, ****p<0.0001 and ns indicates non-significant). Statistical significance determined by Student’s t-test.

We next investigated whether YTHDF2 depletion alters the dynamics of newly synthesised RNA following transcriptional inhibition. Following EU labelling, transcription was inhibited using actinomycin D (25 µg/ml) for 2 hours, and residual EU-associated fluorescence was subsequently quantified. Interestingly, actinomycin D treatment resulted in a marked redistribution of the EU-associated signal from the nucleolar region toward the broader nucleoplasmic compartment **(Supplementary Figure** - **S4C, S4D, S4E)**, consistent with dynamic redistribution and processing of newly synthesised RNA following transcriptional inhibition. To compare the residual EU-associated signal following actinomycin D treatment, the ActD/EU fluorescence ratio was calculated and normalised to the corresponding EV-NG value for each biological replicate. High-glucose treatment resulted in a significant increase in the normalised ActD/EU ratio compared with both NG and HM conditions **(Figure 4E and 4F)**, indicating altered global RNA turnover dynamics under high-glucose stress. Notably, HM treatment reduced residual EU intensity relative to NG, suggesting that hyperosmolarity alone has a distinct effect on global RNA dynamics, whereas HG treatment significantly increased residual EU intensity compared with both NG and HM. The distinct and opposing responses to the osmotic control and high-glucose treatment support the conclusion that the increased ActD/EU ratio observed under HG conditions is attributable predominantly to glucose-associated stress rather than to osmotic effects alone. Comparison of the normalised ActD/EU ratios between EV and YTHDF2-depleted cells revealed a consistent reduction in the ratio across NG, HM, and HG conditions **(Figure 4G)**. This finding indicates that depletion of YTHDF2 alters the dynamics of residual newly synthesised RNA following transcriptional inhibition. Together with the reduction in EU incorporation observed in the pulse-labelling experiment, these findings demonstrate that YTHDF2 depletion perturbs both nascent RNA production and global RNA turnover dynamics. Collectively, these observations suggest that YTHDF2 contributes to the maintenance of global RNA homeostasis by influencing both nascent RNA production and the dynamics of newly synthesised RNA following transcriptional inhibition. Together, our cellular studies suggested that YTHDF2 is an important regulator of renal cell responses to hyperglycemic stress.

We therefore asked whether the reduction in YTHDF2 observed in our experimental models was also evident in human DKD and whether YTHDF2 expression was associated with molecular signatures of renal injury. To address this, we performed a meta-analysis of independent human DKD transcriptomic datasets encompassing distinct renal compartments. Integrating these datasets enabled us to identify gene expression changes that were consistently observed across different patient cohorts, helping distinguish reproducible disease-associated patterns from dataset-specific variations. Furthermore, the inclusion of different renal compartments provided a broader perspective on the molecular alterations associated with DKD and the context-dependent expression of m6A-associated regulators. By examining YTHDF2 expression alongside established inflammatory and fibrotic gene signatures, we sought to determine whether the changes observed in our experimental models were reflected in human disease and to identify potential molecular associations warranting further mechanistic investigation.

### YTHDF2 expression is reduced in human DKD glomeruli and inversely associated with inflammatory and fibrotic gene expression

To determine whether the molecular alterations observed in our experimental models were also observed in human diabetic kidney disease, we performed a targeted analysis of publicly available human DKD transcriptomic datasets representing distinct renal compartments such as GSE142025, comprising kidney biopsy transcriptomes, GSE96804, comprising microdissected glomerular samples and GSE104954, comprising tubulointerstitial transcriptomes. Rather than performing an unrestricted genome-wide comparison, we focused our analysis on predefined gene sets representing inflammatory signaling, oxidative stress, renal fibrosis, and m6A-associated RNA regulatory machinery. This targeted approach was intended to determine whether the molecular processes identified in our experimental models were also represented across independent human DKD cohorts. Notably, the three datasets were generated using different transcriptomic platforms and represented different renal compartments; therefore, concordant alterations observed across these cohorts are particularly informative, while differences between datasets may reflect tissue composition, disease stage, technical platform, or compartment-specific biology.

Analysis of the upregulated genes across these disease-associated categories identified 25 genes that were consistently increased across all three datasets (**Supplementary Figure** - **S5A**), indicating a conserved molecular signature of diabetic renal injury. The combined analysis demonstrated concordant alterations in genes associated with renal fibrotic remodeling, inflammation, oxidative stress, and m6A-associated RNA regulation (**Figure 5A**; individual dataset lollipop plots are shown in **Supplementary Figure** - **S5B, S5C, S5D**). Among the fibrotic genes examined, *COL3A1, MMP2, TIMP1, COL1A1,* and *FN1* showed consistent upregulation across the datasets, whereas *CTGF, LOX,* and *NID1* were reduced. Similarly, inflammatory genes including *C3, VCAM1, CXCL10, CXCL1, NFKB,* and *TLR4* were consistently increased, while *PTGS2* and *IL1B* were reduced. Within the oxidative-stress-associated gene set, *SOD2, NQO1,* and *NFE2L2* were among the genes showing increased expression. Several m6A-associated regulatory factors, including *YTHDC1, IGF2BP3, METTL3, RBM15, YTHDF1,* and *WTAP*, also exhibited increased expression across the analyzed datasets. Collectively, these findings demonstrate that human DKD transcriptomes exhibit conserved alterations in molecular signatures associated with renal fibrosis, inflammation, oxidative stress, and m6A-associated RNA regulation.

**Figure 5:**
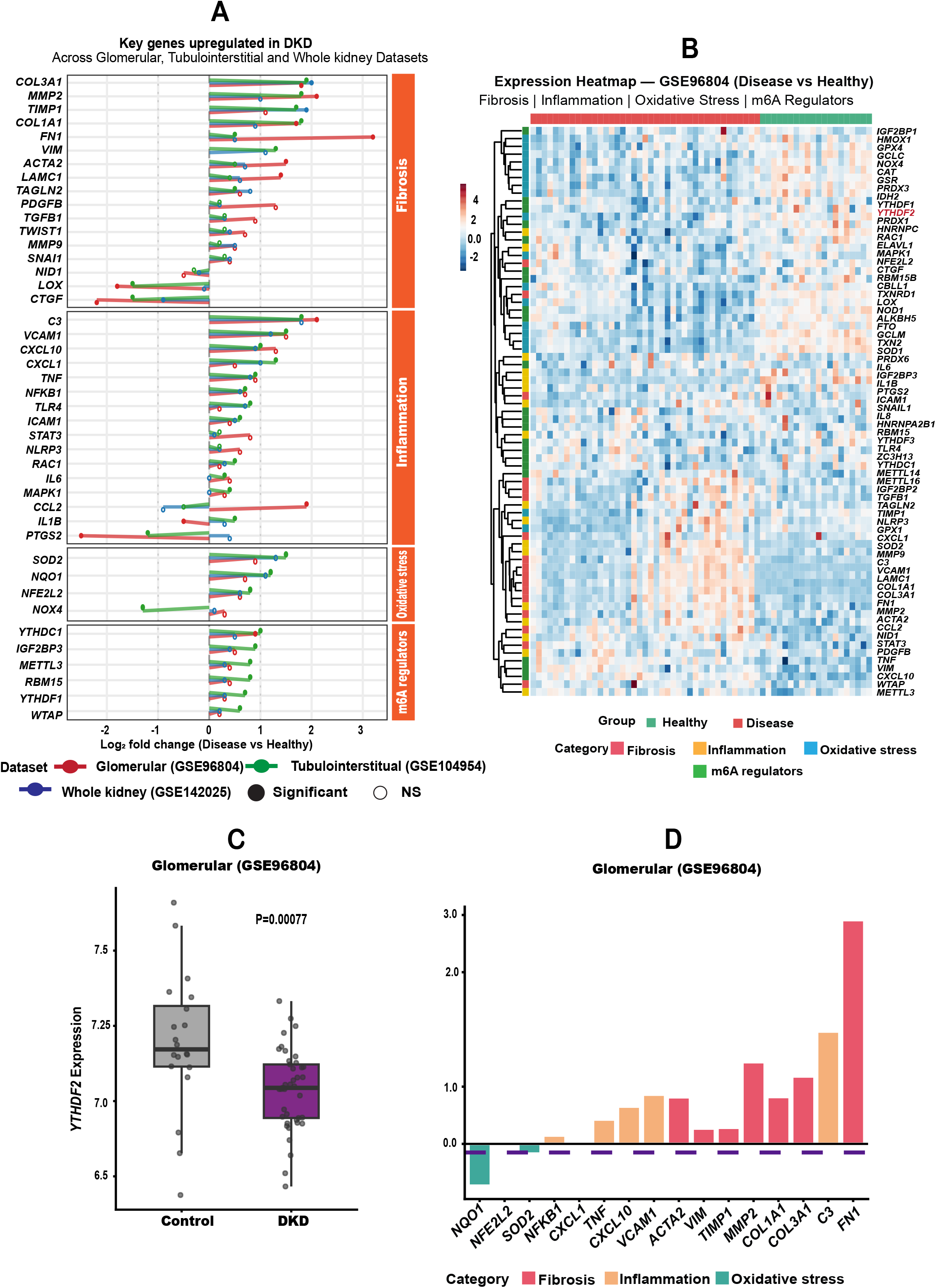
**A.** Log2FC of the upregulated genes plotted per category (fibrosis, inflammation, oxidative stress, and m6A regulators) in diabetic kidney disease (DKD) across three compartment-specific datasets with each dataset represented by a distinct color; filled circles denote significant genes (adj. *p* < 0.05) and open circles indicate non-significant values. **B.** Expression heatmap of key genes across fibrosis, inflammation, oxidative stress, and m6A regulator categories in GSE96804 (glomerular compartment). Each row represents a gene and each column a sample, with color intensity indicating mRNA expression level (RMA-normalized; red = upregulated, blue = downregulated). Samples are annotated by group (Healthy vs. Disease), *YTHDF2* is written in red colour to highlight its expression in control vs healthy. **C.** Box and whisker plot showing *YTHDF2* expression in healthy controls and patients with DKD in the GSE96804 transcriptomic dataset. Each dot represents an individual sample. **D.** Bar plot of log2 fold change (log2FC) (DKD vs Control) for *YTHDF2* and a panel of fibrosis (red), inflammation (orange), and oxidative stress (teal) marker genes in glomerular transcriptomic data from GSE96804. The dashed purple line marks *YTHDF2* log2FC, which is negative, contrasting with the predominantly positive fold changes observed for fibrosis and inflammation markers illustrating an inverse relationship between *YTHDF2* downregulation and upregulation of DKD-associated pathways in the glomerulus.

We next focused on YTHDF2, given its consistent reduction at the transcript level in our HFD+STZ treated mouse kidneys and high-glucose-treated renal cell line models. Within the targeted m6A-associated gene analysis, *YTHDF2* showed a modest reduction in expression in the GSE96804 glomerular dataset **(Figure 5B)**; however, this change did not reach statistical significance in the broader analysis represented in the plot **(Supplementary Figure** - **S5B)**. We therefore performed a focused analysis of YTHDF2 expression in the same dataset, which revealed a significant reduction in YTHDF2 expression in DKD glomerular samples compared with controls **(Figure 5C)**, consistent with the previously reported study (34). Importantly, this finding was concordant with the reduced *Ythdf2* expression observed in diabetic mouse kidney and renal cell hyperglycemic model (Figure 1D, Figure 2C&D), supporting a reproducible association between glucose stress and reduced YTHDF2 expression across different model systems.

Given the established role of YTHDF2 as an m6A reader involved in RNA turnover and decay, we next performed correlation analysis in the GSE96804 glomerular dataset to identify genes inversely associated with YTHDF2 expression, with the aim of identifying candidate transcripts that may be subject to YTHDF2-dependent regulation of RNA stability. YTHDF2 expression exhibited significant inverse associations with multiple genes involved in renal inflammation and fibrotic remodeling, including *FN1, C3, MMP2, COL3A1, COL1A1, TIMP1, TNF, CXCL10, VCAM1, ACTA2,* and *VIM* **(Figure 5D)**. Notably, several of these genes were also consistently upregulated across the independent DKD datasets examined, suggesting that their association with diabetic renal injury represents a conserved molecular signature rather than a dataset-specific observation. The inverse association between YTHDF2 expression and these disease-associated transcripts raises the possibility that YTHDF2-dependent RNA decay may contribute to their post-transcriptional regulation, warranting further investigation into whether these transcripts represent direct targets of YTHDF2-mediated RNA turnover in DKD.

Collectively, these findings extend our observations from experimental models to human DKD and demonstrate that reduced YTHDF2 expression is associated with conserved inflammatory and fibrotic molecular signatures in diabetic kidney disease. Together, these findings support a potential role for YTHDF2 dysregulation in the altered post-transcriptional RNA regulatory landscape associated with DKD.

## Discussion

Diabetic kidney disease (DKD) is characterised by metabolic stress, oxidative stress, inflammation, extracellular matrix remodelling, and progressive renal cellular injury. Growing evidence implicates epitranscriptomic mechanisms, particularly N6-methyladenosine (m6A) RNA regulation, in these processes; however, the contributions of individual m6A-associated regulators to DKD remain incompletely understood. In this study, using HFD+STZ-treated mouse kidney samples and complementary renal cell line models, we demonstrate that hyperglycemic stress is accompanied by remodelling of the m6A regulatory machinery, with a consistent reduction in expression of the m6A reader *Ythdf2*. YTHDF2 depletion in M15 cells altered cellular viability, proliferation, cell cycle progression and oxidative stress response. Analysis of three independent publicly available human DKD transcriptomic datasets further revealed an inverse association between YTHDF2 expression and inflammatory and fibrotic gene signatures in the glomerular compartment. Together, these findings support a potential role for YTHDF2 in regulating post-transcriptional cellular responses associated with diabetic renal injury.

Our targeted transcriptomic and proteomic analyses confirmed prominent inflammatory, oxidative-stress, and fibrotic signatures in HFD+STZ-treated mouse kidney samples, consistent with established molecular features of DKD [44–48]. Differential regulation of podocyte-associated genes further indicated substantial alterations in the podocyte phenotype rather than a uniform loss of podocyte identity, potentially reflecting adaptive or compensatory responses during podocyte stress [49–51]. Importantly, m6A-associated factors were differentially regulated rather than uniformly altered, suggesting remodelling of the m6A regulatory machinery in the diabetic kidney. Notably, the observed upregulation of *Mettl3* together with downregulation of *Mettl14* was consistent with previously reported alterations in these major m6A writer components in diabetic renal injury [29, 30, 52], providing additional support for the remodelling observed in our model. However, *Mettl14* regulation appears to be context dependent, as other studies have reported increased *Mettl14* expression in DKD models, suggesting that its expression and functional contribution may vary with disease stage, renal cell type, or experimental context [31–32].

Interestingly, our analysis also revealed coordinated downregulation of multiple members of the YTH-domain-containing family of m6A readers, including *Ythdc1, Ythdf1, Ythdf2,* and *Ythdf3*. A recent report by Yu et al. (2025) described increased YTHDF2 mRNA and protein expression in renal tissues from their DN model [53]. This variation in the observations, may partly reflect differences in the experimental models, as Yu et al. used a high-fructose diet followed by repeated STZ administration, whereas our study employed a prolonged high-fat diet followed by STZ, warranting studies that investigate the different metabolic stress induced cellular changes. To our knowledge, the simultaneous alteration of these major YTH-domain-containing readers has not previously been characterised in the HFD+STZ diabetic kidney model. These findings raise the possibility that regulation of YTH-domain-containing readers represents an additional layer of m6A-associated RNA regulation during diabetic renal stress. Whether these changes reflect altered upstream mechanisms controlling reader expression, localisation, or functional engagement with m6A-modified transcripts remains to be determined.

Although multiple in vivo models of DKD have been used to investigate the complex molecular and cellular alterations associated with disease progression [54–56], complementary in vitro models remain essential for dissecting cell-intrinsic responses to hyperglycemic stress. In this context, we employed M15 and IMCD3 cells, which represent distinct developmental contexts of the kidney, to characterise the m6A regulatory landscape and its response to glucose stress. The involvement of m6A-associated pathways in kidney development is an emerging area, with recent work by Ramalingam et al. demonstrating that an RNA transmethylation pathway contributes to kidney nephrogenic potential [36]. M15 cells therefore provided an opportunity to examine m6A-associated regulation in an early developmental renal context, while comparison with IMCD3 cells allowed us to determine whether the expression of these regulators varied across distinct developmental contexts. This comparison was not intended as a direct functional comparison between the two cell lines, but rather to establish the cellular context of m6A-associated regulation and identify factors that may be differentially responsive to hyperglycemic stress.

The response of m6A-associated regulators to high-glucose stress was also dependent on renal cellular context. M15 and IMCD3 cells exhibited distinct patterns of m6A-regulator expression following glucose stress, potentially reflecting differences in developmental origin and cell-type-specific signalling programs. These findings extend the emerging concept that metabolic stress can differentially remodel m6A-associated regulatory factors across distinct renal cellular contexts. Among the altered regulators, *Mettl14, Ythdc1,* and *Ythdf2* were consistently reduced in the diabetic kidney and glucose-stressed renal cell models. Previous studies have demonstrated heterogeneous responses of m6A-associated factors to hyperglycemic stress across renal cell types, with METTL14 reported to be downregulated in HG-treated HK-2 cells [48] but upregulated in glomerular endothelial cells [31], whereas YTHDC1 was reported to be downregulated in HG-treated human and mouse podocyte cell lines [57], and YTHDF2 was reported to be transcriptionally upregulated in HG-treated HK-2 cells [53]. These findings further support the context-dependent regulation of individual m6A components across renal cell types and experimental conditions.

Loss of YTHDF2 reduced cellular viability and proliferation while increasing intracellular ROS levels, suggesting that YTHDF2 contributes to the regulation of cellular growth and oxidative-stress responses. Notably, the increase in ROS following YTHDF2 depletion was observed irrespective of glucose condition, suggesting that this effect may reflect a broader role of YTHDF2 in cellular responses to oxidative stress rather than being specific to hyperglycemic stimulation. This observation is consistent with previous studies reporting reduced cellular proliferation following YTHDF2 depletion in other biological contexts [58–60] and with studies implicating YTHDF2 in the regulation of oxidative stress and ROS homeostasis [61–62]. However, the functional consequences of YTHDF2 dysregulation appear to be highly context dependent. For instance, a recent study in renal cell carcinoma reported that YTHDF2 overexpression reduced cellular proliferation, invasion, and migration [63], whereas another study reported reduced ROS levels following YTHDF2 depletion [64]. These apparently divergent findings suggest that the effects of YTHDF2 may vary according to cellular identity, disease context, and the surrounding signalling environment. Our findings extend this context-dependent role of YTHDF2 to an early developmental renal cell model exposed to metabolic stress, in which YTHDF2 depletion was associated with increased oxidative stress and altered cellular growth.

EU-based Click-iT analysis revealed that high-glucose treatment reduced EU incorporation, with a further reduction observed following YTHDF2 depletion, suggesting that YTHDF2 may contribute to nascent RNA-associated activity during hyperglycemic stress. Although YTHDF2 is classically characterised as an m6A reader that promotes cytoplasmic mRNA turnover and decay, our findings raise the possibility that its depletion may also influence processes associated with nascent RNA production or transcription. This interpretation is consistent with the observation by Zhang et al. 2024 that YTHDF2 loss reduced nascent transcript abundance in CD8+ T cells, suggesting that YTHDF2 may have functions extending beyond its canonical role in cytoplasmic mRNA turnover [65]. These findings raise the possibility of a previously underexplored role for YTHDF2 in processes linked to nascent RNA production, although the molecular basis of this effect remains unknown. Whether YTHDF2 directly influences transcription or affects nascent RNA abundance indirectly through its established role in post-transcriptional RNA regulation therefore remains unresolved. The predominant nucleolar EU signal observed following the pulse, together with its redistribution after Actinomycin D treatment, is consistent with a substantial contribution from ribosomal RNA, as previously described [66–68], providing additional support for the observed change in the subcellular distribution of EU-labelled RNA following transcriptional inhibition.

Based on the established role of YTHDF2 in promoting m6A-dependent RNA turnover, we initially expected YTHDF2 depletion to result in increased retention of labelled RNA following transcriptional inhibition. However, contrary to this expectation, YTHDF2-depleted cells exhibited reduced residual EU signal following Actinomycin D treatment (**Figure 4C**, **4E and 4G**). This apparent variation should be interpreted in the context of the characteristics and limitations of the EU-based assay. One possible explanation is the relatively low EU incorporation observed in YTHDF2-depleted cells following the initial pulse, resulting in a smaller pool of labelled RNA available for subsequent detection after transcriptional inhibition. Thus, although the Click-iT EU/Actinomycin D approach provides an overall measure of labelled RNA retention, it cannot be used to directly infer mRNA turnover or YTHDF2-dependent mRNA stability, particularly given the substantial contribution of rRNA to the measured EU signal. Nevertheless, the differential residual EU signal following transcriptional inhibition indicates that YTHDF2 depletion is associated with altered cellular RNA dynamics or RNA homeostasis. These findings therefore raise the possibility that YTHDF2 contributes to RNA homeostasis in renal cells under metabolic stress, although the precise processes involved remain unresolved. Transcript-specific approaches, such as Actinomycin D chase followed by RT-qPCR, together with assessment of YTHDF2–RNA association and m6A modification, will be required to determine whether YTHDF2 depletion alters the stability or turnover of specific mRNAs and to distinguish effects on RNA stability from changes in transcription, RNA processing, or other components of RNA metabolism.

Analysis of three publicly available human DKD transcriptomic datasets provided further relevance to our experimental findings, with YTHDF2 significantly downregulated in the glomerular dataset, consistent with the reduction observed in our experimental models. In contrast, YTHDC1 was increased across the three human datasets, differing from the downregulation observed in our experimental models and further highlighting the context-dependent regulation of individual m6A-associated factors. The inverse association of YTHDF2 with inflammatory and fibrotic transcripts in glomeruli raises the possibility that YTHDF2 may influence their RNA stability or turnover. Further investigation is required to establish whether these transcripts are direct YTHDF2 targets and whether their regulation occurs through m6A-dependent mechanisms.

Given the context-dependent regulation of m6A-associated factors across renal cell types under high glucose in our study and previous reports [31, 48, 53, 57], the functional relevance of YTHDF2 identified in M15 cells warrants further validation in additional DKD-relevant renal cell types, particularly podocytes, tubular epithelial cells, and glomerular endothelial cells, as well as in appropriate in vivo models. Furthermore, the direct RNA targets and mechanisms through which YTHDF2 may regulate inflammatory and fibrotic responses, including its potential effects on RNA stability or turnover, remain to be established. Validation of these mechanisms and their association with disease stage in clinically characterised patient cohorts will be important to determine the broader and translational relevance of YTHDF2-regulated transcripts as biomarkers in DKD. Given the limitations of current markers such as albuminuria and eGFR in capturing early or progressive renal injury [69], identifying complementary biomarkers may improve the detection and stratification of DKD.

Overall, our findings identify YTHDF2 as a potential regulator of renal cellular responses to metabolic stress and provide evidence linking its altered expression to cellular dysfunction and disease-associated molecular changes in DKD. These findings highlight m6A-associated post-transcriptional regulation as a potential component of renal adaptation to diabetic stress.

## Acknowledgements

We thank DBT (BT/RLF/Re-entry/21/2019), IISER Berhampur seed grant and PMRF for the funding. We thank Dr. Shovamayee Maharana and Prerna Nawal, MCB, IISc for help with the click chemistry experiment. We extend our thanks to IISER Berhampur CAIF, imaging facility and Flow Cytometry facility. We would like to thank Nishad Gosavi, RBL, DBS, IISER Berhampur for their support.

## Author contributions

H.S.S.H. and R.S.B. conceived and designed the study. H.S.S.H. performed the in vitro experiments, RT-qPCR, generation and characterization of genome-edited cell lines, and data analysis. A.K.P. performed in vitro western blotting experiments and analyzed the data. K.T. performed the meta-analysis and corresponding data visualization. N.S. and A.K. performed the animal experiments. V.K.P. provided resources and support for the animal experiments. H.S.S.H. drafted the manuscript. R.S.B. supervised the study, acquired funding, and revised the manuscript. All authors reviewed and approved the final manuscript.

