## Supporting Information for "The RNA m6A reader YTHDF2 depletion phenocopies high glucose induced-cellular stress response in diabetic kidney disease"

**Supplementary Table 1:** Guide RNA sequences and/or oligonucleotide sequences used for generation of YTHDF2-depleted M15 cells.

**Supplementary Table 2:** Primer sequences used for RT-qPCR analysis.

**Supplementary Table 3:** List of genes included in the targeted meta-analysis of inflammatory, oxidative stress, fibrotic, and m6A-associated markers.

**Figure Legends**

**Supplementary Figure S1. Additional characterization of the HFD+STZ-induced DKD mouse model**

**A.** Gene ontology (GO) enrichment analysis of the transcriptomic dataset comparing diabetic and healthy kidney samples. The dot plot represents the top enriched biological processes, with dot size indicating gene count and color indicating adjusted *p*-value. **B.** Lollipop plot depicting significantly differentially expressed proteins identified by proteomic analysis of diabetic versus healthy kidney samples. Proteins are grouped according to their predominant biological association with inflammation, oxidative stress, cellular stress, and podocyte injury. **C.** GO enrichment analysis of the proteomic dataset showing the top enriched biological processes in diabetic versus healthy kidney samples. Dot size represents gene/protein count, and color indicates adjusted *p*-value. **D.** Heatmap showing ΔCt values of m6A-associated regulatory factors, including writers, erasers, and readers, in control and HFD+STZ-treated kidney samples.

**Supplementary Figure S2. Characterization of m6A-associated regulatory factors in renal cell models and their response to high-glucose stress**

**A.** Heat map representing delta Ct values of m6A modifier expression in the early mesonephros-derived M15 cell line in comparison with the metanephros-derived inner medullary collecting duct cell line

(IMCD3). **B.** Bar plot displaying relative mRNA expression of significantly dysregulated m6A writers and readers in M15 compared to IMCD3. **C.** Heat map representing delta Ct values of m6A regulators in high-glucose (HG -30mM) treated IMCD3 cell lines with respect to the normal-glucose-treated cell line (NG-5.5 mM). **D.** Relative gene expression of Ythdf2 in HG-treated M15 cells in comparison with NG and HM-treated M15 cells.

**Supplementary Figure S3. Additional characterization of YTHDF2-depleted M15 cells.**
**A.** Schematic of guide RNA (Sg1 and Sg2) target sites in YTHDF2. **B.** Bright-field image indicating genome-edited M15 cells showing a more elongated cell appearance (mesenchymal-like) in KD1 and KD2 compared to EV. **C.** Scatter plot indicating elevated aspect ratio of YTHDF2-transduced cells with respect to EV (analysis from bright-field images), suggesting a more mesenchymal nature of YTHDF2 KD cells irrespective of treatment conditions (NG, HM and HG, n=3*50 cells). **D.** Scatter plot indicating decreased solidity of YTHDF2 guide RNA-transduced cells with respect to EV (analysis from bright-field images), suggesting a more mesenchymal nature of YTHDF2 KD cells irrespective of treatment conditions (NG, HM and HG, n=3*50 cells). Data are presented as a scatter plot showing the distribution of individual observations. **E.** Histogram indicating PI-stained cells in different stages of the cell cycle (G1/G0, S, and M phase), plotted for YTHDF2 KD cells (KD1 and KD2) with respect to M15 EV. **F.** Bar graphs indicating the % cells of YTHDF2 KD cells with respect to M15 EV cells, representing cell cycle stages such as G1/G0, S, and G2/M. **G.** Bar graph representing the increased % cells in S phase in YTHDF2 KD cells compared to control cells, suggesting S phase arrest of the YTHDF2 KD cells in the cell cycle. **H.** Bar plot indicating validation of ROS production in DCFDA-stained cells with respect to DCFDA-unstained cells (NC) in YTHDF2 KD cells with respect to EV. PC (Positive control) represents H2O2-treated DCFDA-stained cells showing elevated ROS compared to control EV cells (EV NG). Data are presented as mean ± SD from $N$ = 3 independent biological replicates. Asterisks indicate statistical significance. (*p < 0.05, * *p < 0.01, * * *p < 0.001 and * * * *p < 0.0001 ) determined by Student's t-test)

**Supplementary Figure S4. Image representing Eu-labelled click chemistry**
**A.** Representative image indicating negative control (cells incubated with just media and stained using AF-577 in EV, KD1 and KD2). **B.** Representative image indicating another negative control (cells first treated with Act D followed by incubation of cells with EU and stained using AF-577). Representative image showing nascent RNA synthesis measured by EU (5-Ethynyl uridine) staining and EU-labelled cells after treatment with Actinomycin-D for 2 hours in **C.** empty vector control, **D.** YTHDF2 KD1, and **E.** YTHDF2 KD2 across NG, HM, and HG (DAPI represents nuclei stained in blue, AF-577 indicates intensity of EU in red, and Composite indicates overlay of DAPI and AF-577).

**Supplementary Figure S5. Targeted analysis of human DKD transcriptomic datasets**
**A.** Venn diagram showing the overlapping regions and the number of shared and unique upregulated genes among glomerular (GSE96804), tubulointerstitial (GSE104954), and whole kidney (GSE142025) datasets, with 25 genes commonly upregulated across all three. Lollipop plot showing $\log_2$ fold change of key genes (Disease vs. Healthy) across four gene categories such as fibrosis, inflammation, oxidative stress, and m6A regulators in **B.** GSE96804, **C.** GSE142025, and **D.** GSE104954. Filled circles in the lollipop plot denote statistically significant genes (adj. $p < 0.05$, $|\log_2 FC| \geq 1$); open circles indicate non-significant or subthreshold genes

**Supplementary Table 1:**

| Genes | Single guide RNA sequence (5'-3') | Targeting exons |
|---|---|---|
| YTHDF2 | Sg1- GGCATGAATACTATAGACCAAGG | 4 |
| | Sg2- CCACGACCTTGACGTTCCTAAGG | 5 |

**Supplementary Table 2:**

| | Gene | Forward Primer (5'-3') | Reverse primer (5'-3') |
|---|---|---|---|
| 1 | mMETTL3 | **CCGTAGTGATAGTCCCGTGC** | **GCAAGACGGATGGAAACAGC** |
| 2 | mMETTL14 | **GACTGGCATCACTGCGAATG** | **AGGTCCAATCCTTCCCCAGA** |
| 3 | mVIRMA | **TCTTTCCCCACAATAGCCCG** | **GCTGTGAAGATACCTAAAGAGGACA** |
| 4 | mRBM15 | **CGCTTCTAGGCTTTGGGTCA** | **ACGCCCTAATCCCAGTACCT** |
| 5 | mRBM15B | **CGTGGGAGGACAACTCACTC** | **GAGTTGCTGCCTGACTCCTT** |
| 6 | mWTAP | **CATTTTGTGGCTGCGAGACC** | **ACTCATCCCGTGCCATAACT** |
| 7 | mHAKAI | **CTGACTTCCGTTCCCACTGT** | **AATCCTTCATCACCTTGCGGG** |
| 8 | mZC3H13 | **CTTCTGTGCGTCTCTCGTCT** | **CCCTTGTAAGCGCAGTCTTCA** |
| 9 | mFTO | **GAGGAGCCCTATTTCGGCAT** | **GGTGTCTCGATGTCCCAAGA** |
| 10 | mALKBH5 | **TGTCGGAACCTGTGCTTTCT** | **GAGGCCGTATGCAGTGAGTG** |
| 11 | mYTHDC1 | **GGAAGAGAACCAGAGCACGG** | **GTTCCCCATCTTTCTCCTCCC** |
| 12 | mYTHDF1 | **CAACCTACTTCTGCCCCGAG** | **TTGATGGAGCGGTGGATGTC** |
| 13 | mYTHDF2 | **CAGGCATCAGTAGGGCAACA** | **TAGATCCAGAACCCGCCTGA** |
| 14 | mYTHDF3 | **GGATCAGAGACCTAAAGGGCAA** | **CCATGCTGCTTCCCCAAGAG** |
| 15 | mHNRNPA2B1 | **GTCCGCGATGGAGAGAGAAA** | **TCCCGCATAACCACACAGTC** |
| 16 | mHNRNPG | **CAGATCGCCCAGGAAAGCTC** | **TCTGCTGGGCTTTCGAGTGTA** |
| 17 | mIGF2BP1 | **GGCTCACCCAAACTCTCCC** | **CTCTCGTTGAGGTTGCCGAT** |
| 18 | mIGF2BP2 | **ACCAAACGCCAGACGAGAAT** | **CCTGATCTTGCGTTGTGCAG** |
| 19 | mIGF2BP3 | **TCAAAAAGCCGGACACTCGG** | **TCTTCCTCCCCAACCCCTTT** |
| 20 | mHUR | **CCGCCTCCTTTTCCTGACTT** | **TCTGCACCCTCAAAGCAGAG** |

| | | | |
|---|---|---|---|
| 21 | mPODOCIN | **CACGGGAGAAGTAGCAGGTC** | **GTCCACGTCCACTACGGTG** |
| 22 | mWT1 | **CATCCCAGGCAGGAAAGTGT** | **CTCCTTCCGGCAAACCTGAT** |
| 23 | mPODOCALYXIN | **GGATTTACACCTCCCCTGCC** | **AACCTTTCCCCGTGACCATC** |
| 24 | mSYNAPTOPODIN | **CGTAGCCAGGTGAGCCAAG** | **GTTGAAGAGCTGGACCCCTC** |
| 25 | mNEPHRIN | **AAAGCAGGGCACACAGAGAG** | **CCTTCCACCACAGTCAGGTT** |
| 26 | mALPHA ACTININ 4 | **TGGTGCCAGAGAAAGACAGC** | **CGATGTCCTCAGCATCCAAC** |
| 27 | mCD2AP | **TTGAATGGCCTTGTGCTGTAG** | **GAAGGCAAGGTAGGCTAAACG** |
| 28 | mTRPC6 | **CTGTCTCCCAAGCCCCTAAC** | **CCTGACTGGCTAGGTTCTGC** |
| 29 | mGAPDH | **AACTTTGGCATTGTGGAAGG** | **GGATGCAGGGATGATGATGTTCT** |
| 31 | m18SrRNA | **GTAACCCGTTGAACCCCATT** | **CCATCCAATCGGTAGTAGCG** |

**Supplementary Table 3:**

| Sl. No. | Category | Genes |
|---|---|---|
| 1 | **Fibrosis-associated genes** | *Col1a1, Col1a2, Col3a1, Ccn2, Cdh1, Cdh11, Cdh13, Cdh16, Cdh2, Cdh3, Cdh4, Cdh5, Cdh6, Cdhr2, Cdhr5, Vim, Timp2, Timp3, Mmp11, Mmp12, Mmp13, Mmp14, Mmp23, Snai2, Zeb1, Zeb2, Acta2, Tagln, Tagln2, Pdgfrb* |
| 2 | **Oxidative stress-associated genes** | *Hmox1, Hmox2, Nfe2l1, Nfe2l2, Txnip, Nox4, Cyba, Cybb, Nqo1, Gclc, Gclm, Cycs, Apaf1, Coq2, Coq6* |
| 3 | **m6A modulators** | *Mettl17, Mettl1, Mettl16, Mettl2, Mettl6, Mettl3, Mettl7b, Mettl26, Mettl21a, Mettl14, Mettl23, Mettl7a3, Mettl15, Mettl7a2, Mettl4, Mettl15, Mettl7a1, Mettl5, Mettl8, Mettl18, Mettl27, Mettl22, Mettl25, Mettl9, Wtap, Virma, Rbm15, Rbm15b, Zcchc4, Cbll1 (hakai), Wt1, Ythdc2, Ythdc1, Ythdf1, Ythdf2, Ythdf3, Igf2bp1, Igf2bp2, Hnrnpc, Hnrnpa2b1, Rbmx (HNRNPG), Eif3a, Prrc2a, Fto, Alkbh5* |
| 4 | **Apoptosis-associated genes** | *Casp8, Casp6, Casp8ap2, Casp7, Casp9, Casp2, Casp3, Casp4, Casp12, Casp1, Bax, Bak1, Bcl2, Bcl2l12, Bcl2l1, Bcl2l13, Bcl2l11, Bcl2l2, Bcl2a1d, Mcl1, Trp53, Trp53bp2, Trp53inp1, Trp53inp2, Trp53rka, Trp53rkb, Trp53bp1, Trp53i11, Cdkn1a, Pmaip1, Xbp1, Mtor, Akt1* |

| | | |
|---|---|---|
| 5 | **Inflammation-associated genes** | *Tgfb1, Tgfbr1, Tgfbr3, Tgfb1i1, Tgfbr2, Tgfbi, Tgfb2, Tgfbrap1, Ilf2, Il16, Il17ra, Il5ra, Il17rb, Il13ra1, Il6st, Il1rap, Ildr1, Il10rb, Il15ra, Il33, Il18r1, Il1r1, Il6ra, Il17rc, Il4ra, Ilk, Il15, Il34, Il10ra, Ilf3, Ilvbl, Il18, Il7, Ildr2, Il17rd, Il17re, Ilrun, Il18bp, Il11ra1, Il11ra2, Nfkbia, Nfkbie, Nfkb2, Nfkb1, Nfkbib, Nfkbiz, Nfkbil1, C1qtnf1, Tnfaip1, Tnfaip3, Tnfaip2, Tnfrsf10b, C1qtnf12, Tnfrsf12a, Tnfrsf21, Tnfrsf11a, Tnfrsf1b, Tnfrsf1a, Tnfsf13b, Tnfsf10, C1qtnf2, Tnfsf15, C1qtnf3, C1qtnf7, Tnfaip8, Tnfsf12, Inhbb, Inhba, Bmp2, Bmp2k, Bmp4, Bmp6, Bmp7, Acvr2a, Bmpr2, Acvr2b, Acvr1b, Smad5, Smad4, Smad2, Smad7, Smad9, Smad1, Smad3, Smad6, Ccl19, Ccl19-ps1, Ccl19-ps3, Ccl21a, Ccl21b, Ccl21d, Ccl25, Ccl27a, Ccl28, Ccl5, Ccl6, Ccl9, Cxcl10, Ddit3, Rela, Nlrp6* |
| 6 | **Cytoskeletal instability-associated genes** | *Synpo, Synpo2, Nphs1, Nphs2, Podxl, Cd2ap, Actn4, Inf2, Arhgdia, Arhgap24, Rap1gap2, Itgb4, Lamb2, Itga3* |

S 1

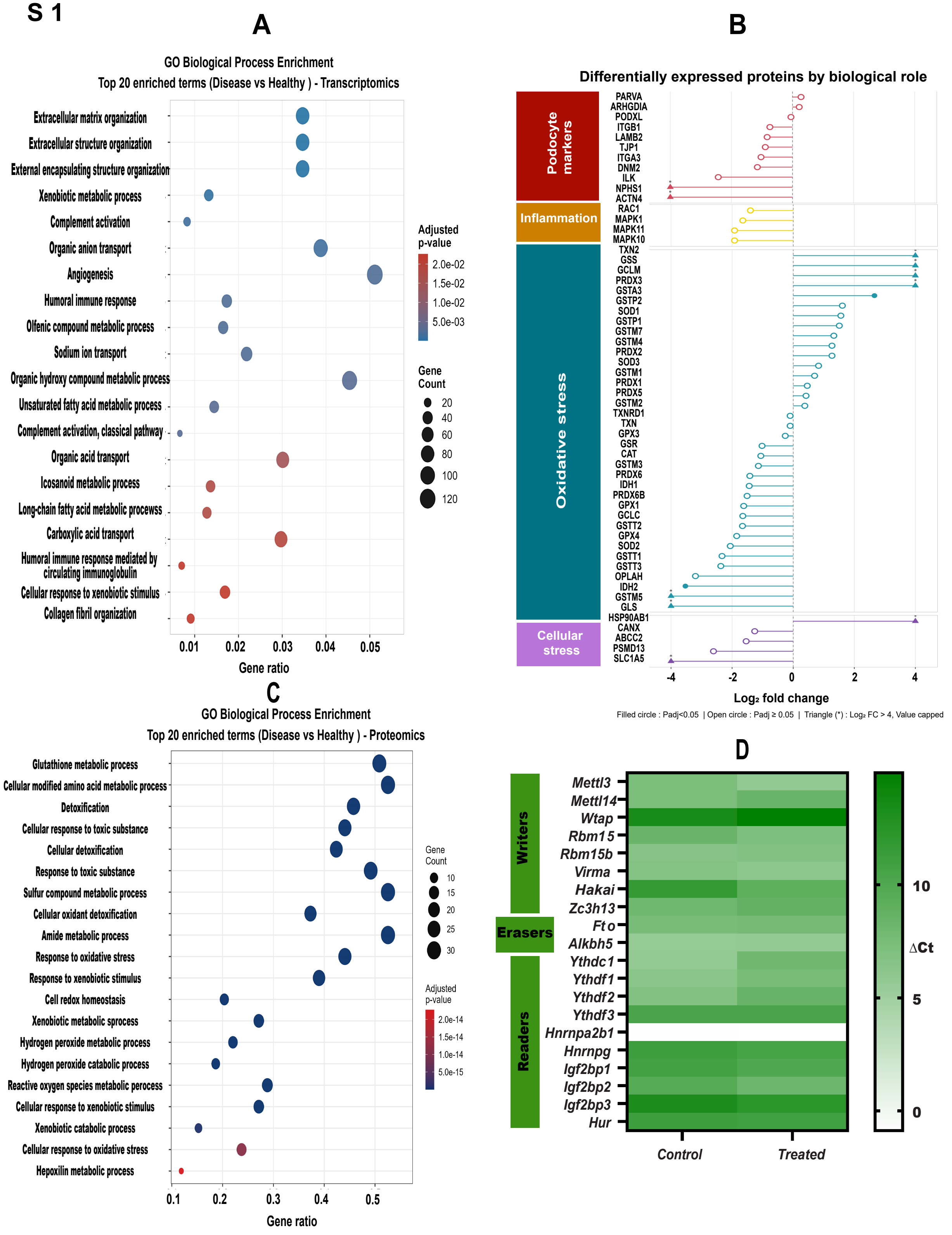
A
GO Biological Process Enrichment
Top 20 enriched terms (Disease vs Healthy ) - Transcriptomics
Extracellular matrix organization
Extracellular structure organization
External encapsulating structure organization
Xenobiotic metabolic process
Complement activation
Organic anion transport
Angiogenesis
Humoral immune response
Olfenic compound metabolic process
Sodium ion transport
Organic hydroxy compound metabolic process
Unsaturated fatty acid metabolic process
Complement activation, classical pathway
Organic acid transport
Icosanoid metabolic process
Long-chain fatty acid metabolic procewss
Carboxylic acid transport
Humoral immune response mediated by circulating immunoglobulin
Cellular response to xenobiotic stimulus
Collagen fibril organization
0.01
0.02
0.03
0.04
0.05
Gene ratio
Adjusted p-value
2.0e-02
1.5e-02
1.0e-02
5.0e-03
Gene Count
20
40
60
80
100
120
B
Differentially expressed proteins by biological role
Podocyte markers
Inflammation
Oxidative stress
Cellular stress
PARVA
ARHGDIA
PODXL
ITGB1
LAMB2
TJP1
ITGA3
DNM2
ILK
NPHS1
ACTN4
RAC1
MAPK1
MAPK11
MAPK10
TXN2
GSS
GCLM
PRDX3
GSTA3
GSTP2
SOD1
GSTP1
GSTM7
GSTM4
PRDX2
SOD3
GSTM1
PRDX1
PRDX5
GSTM2
TXNRD1
TXN
GPX3
GSR
CAT
GSTM3
PRDX6
IDH1
PRDX6B
GPX1
GCLC
GSTT2
GPX4
SOD2
GSTT1
GSTT3
OPLAH
IDH2
GSTM5
GLS
HSP90AB1
CANX
ABCC2
PSMD13
SLC1A5
-4
-2
0
2
4
Log₂ fold change
Filled circle : Padj<0.05 | Open circle : Padj ≥ 0.05 | Triangle (*) : Log₂ FC > 4, Value capped
C
GO Biological Process Enrichment
Top 20 enriched terms (Disease vs Healthy ) - Proteomics
Glutathione metabolic process
Cellular modified amino acid metabolic process
Detoxification
Cellular response to toxic substance
Cellular detoxification
Response to toxic substance
Sulfur compound metabolic process
Cellular oxidant detoxification
Amide metabolic process
Response to oxidative stress
Response to xenobiotic stimulus
Cell redox homeostasis
Xenobiotic metabolic sprocess
Hydrogen peroxide metabolic process
Hydrogen peroxide catabolic process
Reactive oxygen species metabolic perocess
Cellular response to xenobiotic stimulus
Xenobiotic catabolic process
Cellular response to oxidative stress
Hepoxilin metabolic process
0.1
0.2
0.3
0.4
0.5
Gene ratio
Gene Count
10
15
20
25
30
Adjusted p-value
2.0e-14
1.5e-14
1.0e-14
5.0e-15
D
Writers
Erasers
Readers
Mettl3
Mettl14
Wtap
Rbm15
Rbm15b
Virma
Hakai
Zc3h13
Fto
Alkbh5
Ythdc1
Ythdf1
Ythdf2
Ythdf3
Hnrnpa2b1
Hnrnpg
Igf2bp1
Igf2bp2
Igf2bp3
Hur
Control
Treated
10
ΔCt
5
0

S 2

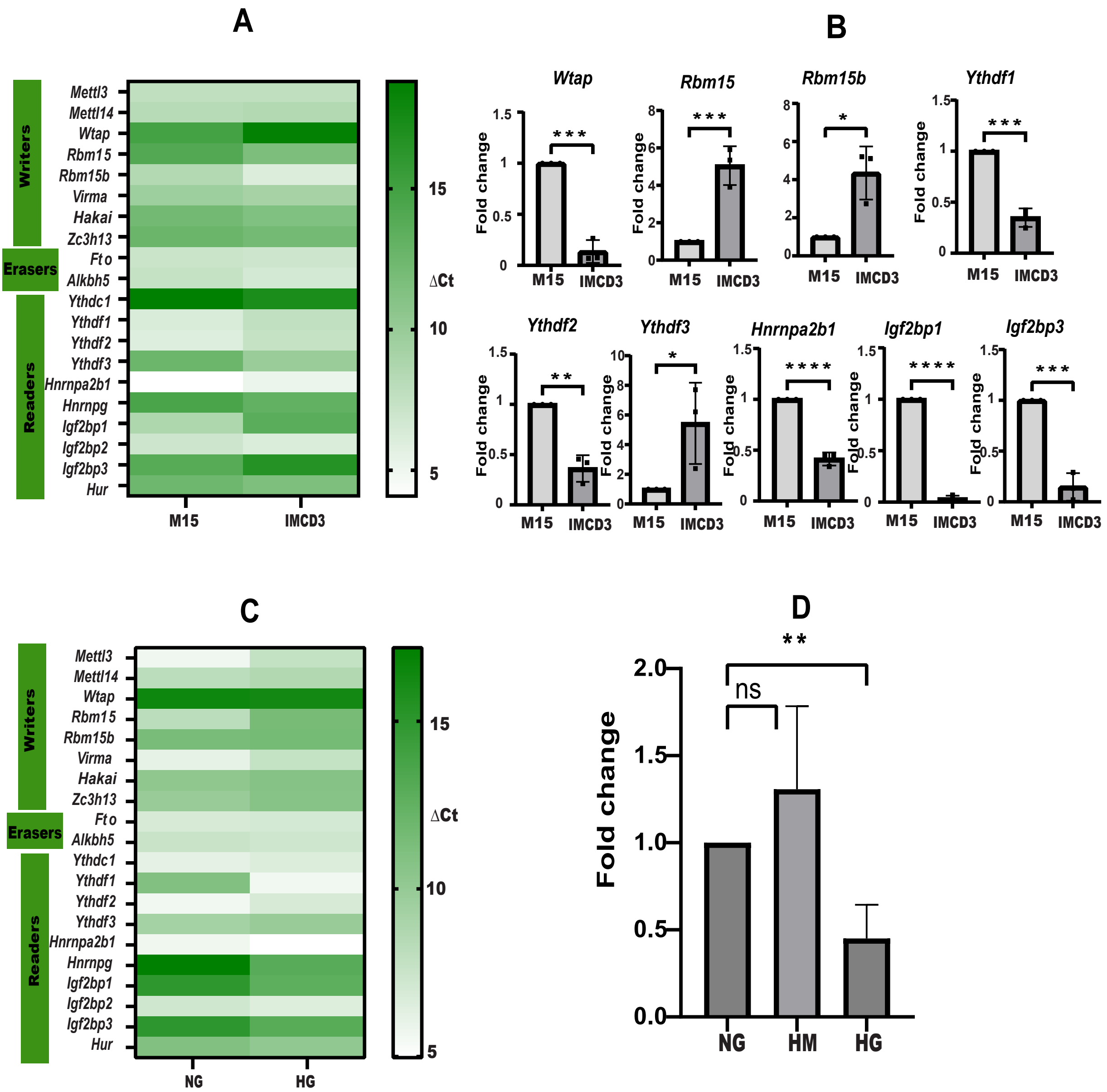
A
Writers
Erasers
Readers
Mettl3
Mettl14
Wtap
Rbm15
Rbm15b
Virma
Hakai
Zc3h13
Fto
Alkbh5
Ythdc1
Ythdf1
Ythdf2
Ythdf3
Hnrnpa2b1
Hnrnpg
Igf2bp1
Igf2bp2
Igf2bp3
Hur
M15
IMCD3
15
10
5
ΔCt
B
Wtap
Rbm15
Rbm15b
Ythdf1
Ythdf2
Ythdf3
Hnrnpa2b1
Igf2bp1
Igf2bp3
Fold change
M15
IMCD3
C
NG
HG
D
ns
**
Fold change
NG
HM
HG

S 3

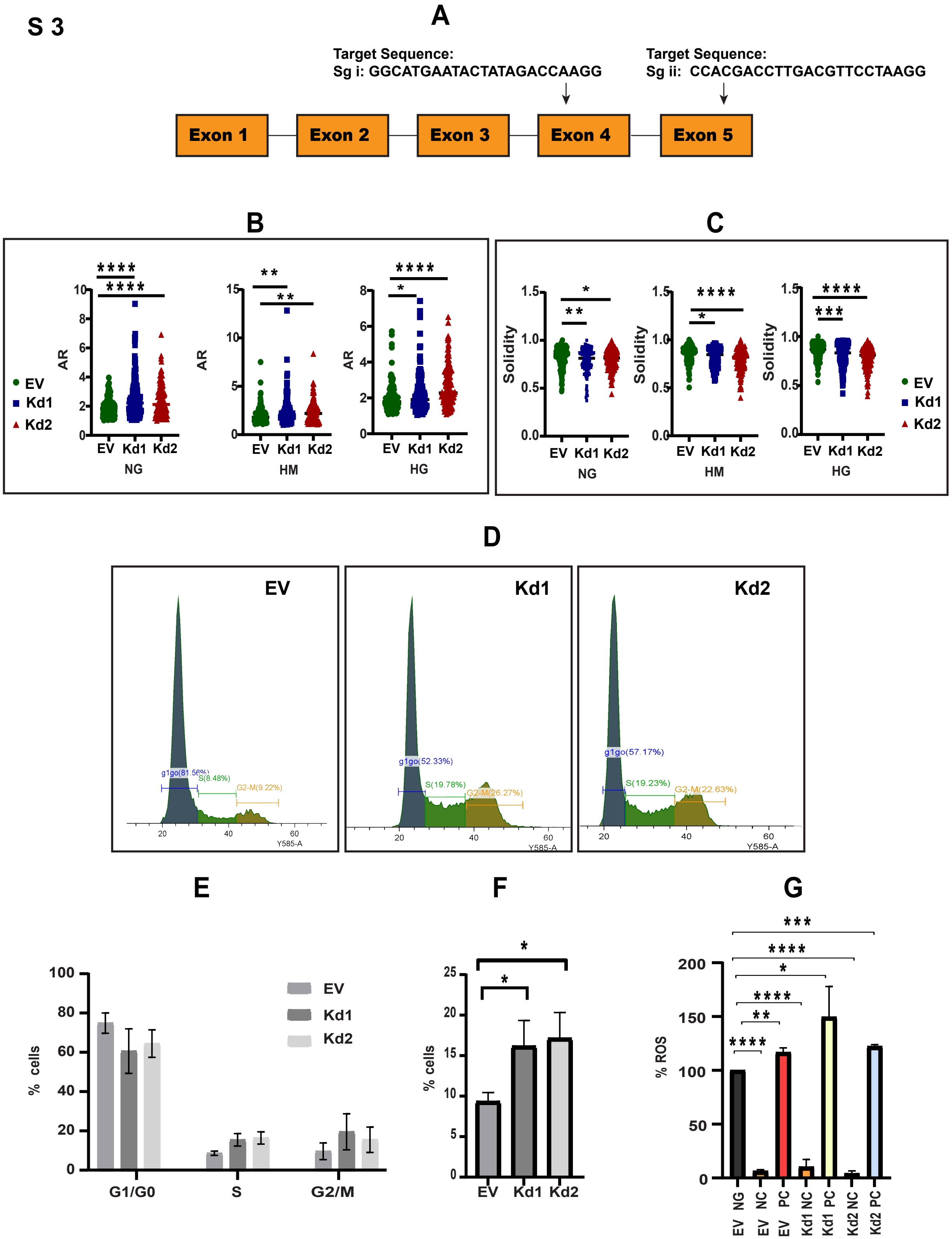
A
Target Sequence:
Sg i: GGCATGAATACTATAGACCAAGG
Target Sequence:
Sg ii: CCACGACCTTGACGTTCCTAAGG
Exon 1
Exon 2
Exon 3
Exon 4
Exon 5
B
AR
EV
Kd1
Kd2
NG
HM
HG
C
Solidity
D
EV
Kd1
Kd2
g1go(81.56%)
S(8.48%)
G2-M(9.22%)
g1go(52.33%)
S(19.78%)
G2-M(26.27%)
g1go(57.17%)
S(19.23%)
G2-M(22.63%)
Y585-A
E
% cells
G1/G0
S
G2/M
F
G
% ROS
EV NG
EV NC
EV PC
Kd1 NC
Kd1 PC
Kd2 NC
Kd2 PC

S 4

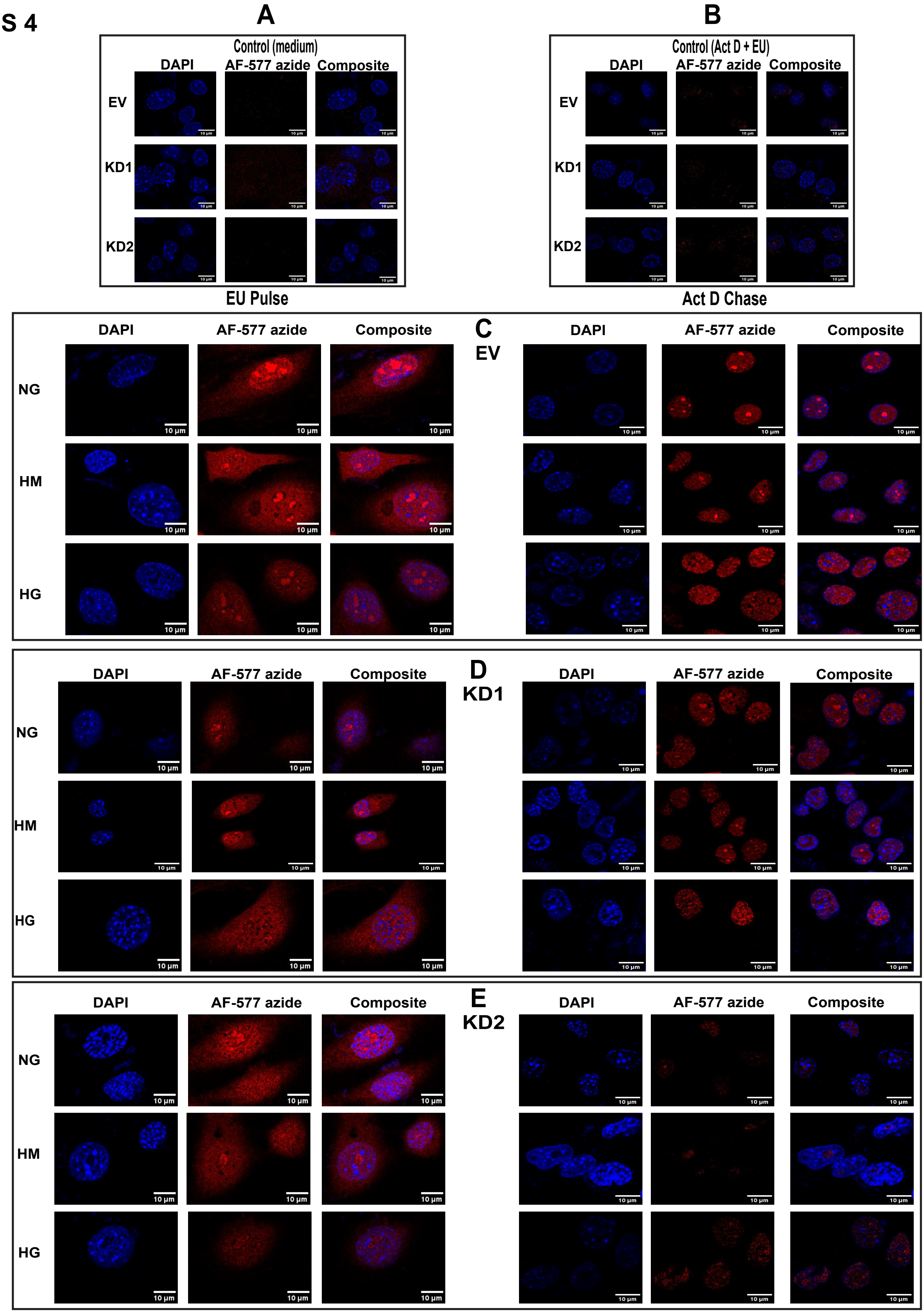
A
B
Control (medium)
Control (Act D + EU)
DAPI
AF-577 azide
Composite
EV
KD1
KD2
EU Pulse
Act D Chase
C
EV
D
KD1
E
KD2
NG
HM
HG
10 µm

S 5

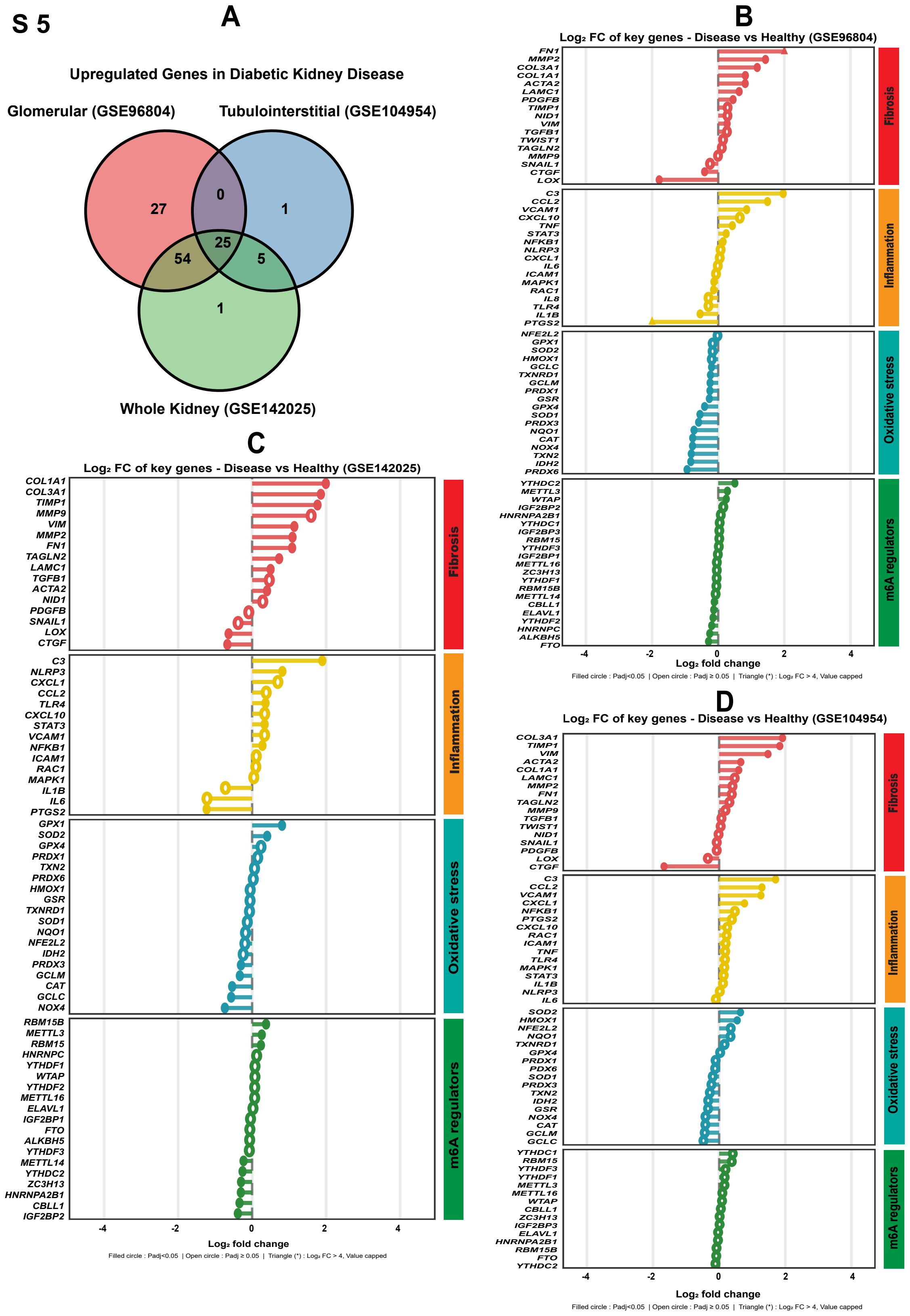
A
Upregulated Genes in Diabetic Kidney Disease
Glomerular (GSE96804)
Tubulointerstitial (GSE104954)
Whole Kidney (GSE142025)
27
0
1
54
25
5
1
B
Log2 FC of key genes - Disease vs Healthy (GSE96804)
FN1 MMP2 COL3A1 COL1A1 ACTA2 LAMC1 PDGFB TIMP1 NID1 VIM TGFB1 TWIST1 TAGLN2 MMP9 SNAIL1 CTGF LOX
Fibrosis
C3 CCL2 VCAM1 CXCL10 TNF STAT3 NFKB1 NLRP3 CXCL1 IL6 ICAM1 MAPK1 RAC1 IL8 TLR4 IL1B PTGS2
Inflammation
NFE2L2 GPX1 SOD2 HMOX1 GCLC TXNRD1 GCLM PRDX1 GSR GPX4 SOD1 PRDX3 NQO1 CAT NOX4 TXN2 IDH2 PRDX6
Oxidative stress
YTHDC2 METTL3 WTAP IGF2BP2 HNRNPA2B1 YTHDC1 IGF2BP3 RBM15 YTHDF3 IGF2BP1 METTL16 ZC3H13 YTHDF1 RBM15B METTL14 CBLL1 ELAVL1 YTHDF2 HNRNPC ALKBH5 FTO
m6A regulators
-4 -2 0 2 4
Log2 fold change
Filled circle : Padj<0.05 | Open circle : Padj ≥ 0.05 | Triangle (*) : Log2 FC > 4, Value capped
C
Log2 FC of key genes - Disease vs Healthy (GSE142025)
COL1A1 COL3A1 TIMP1 MMP9 VIM MMP2 FN1 TAGLN2 LAMC1 TGFB1 ACTA2 NID1 PDGFB SNAIL1 LOX CTGF
Fibrosis
C3 NLRP3 CXCL1 CCL2 TLR4 CXCL10 STAT3 VCAM1 NFKB1 ICAM1 RAC1 MAPK1 IL1B IL6 PTGS2
Inflammation
GPX1 SOD2 GPX4 PRDX1 TXN2 PRDX6 HMOX1 GSR TXNRD1 SOD1 NQO1 NFE2L2 IDH2 PRDX3 GCLM CAT GCLC NOX4
Oxidative stress
RBM15B METTL3 RBM15 HNRNPC YTHDF1 WTAP YTHDF2 METTL16 ELAVL1 IGF2BP1 FTO ALKBH5 YTHDF3 METTL14 YTHDC2 ZC3H13 HNRNPA2B1 CBLL1 IGF2BP2
m6A regulators
-4 -2 0 2 4
Log2 fold change
Filled circle : Padj<0.05 | Open circle : Padj ≥ 0.05 | Triangle (*) : Log2 FC > 4, Value capped
D
Log2 FC of key genes - Disease vs Healthy (GSE104954)
COL3A1 TIMP1 VIM ACTA2 COL1A1 LAMC1 MMP2 FN1 TAGLN2 MMP9 TGFB1 TWIST1 NID1 SNAIL1 PDGFB LOX CTGF
Fibrosis
C3 CCL2 VCAM1 CXCL1 NFKB1 PTGS2 CXCL10 RAC1 ICAM1 TNF TLR4 MAPK1 STAT3 IL1B NLRP3 IL6
Inflammation
SOD2 HMOX1 NFE2L2 NQO1 TXNRD1 GPX4 PRDX1 PDX6 SOD1 PRDX3 TXN2 IDH2 GSR NOX4 CAT GCLM GCLC
Oxidative stress
YTHDC1 RBM15 YTHDF3 YTHDF1 METTL3 METTL16 WTAP CBLL1 ZC3H13 IGF2BP3 ELAVL1 HNRNPA2B1 RBM15B FTO YTHDC2
m6A regulators
-4 -2 0 2 4
Log2 fold change
Filled circle : Padj<0.05 | Open circle : Padj ≥ 0.05 | Triangle (*) : Log2 FC > 4, Value capped
